# Spatial Regulation of CAR Signaling Enables Logic-Gated Activity

**DOI:** 10.64898/2026.05.22.726983

**Authors:** Tushar D. Nichakawade, Brian J. Mog, Alexander H. Pearlman, Taha S. Ahmedna, Nikita Marcou, Ricardo Mora-Cartin, Evangeline Watson, Bum Seok Lee, Sarah R. DiNapoli, Nickolas Papadopoulos, Chetan Bettegowda, Marco Dal Molin, Denis Wirtz, Drew M. Pardoll, Suman Paul, Surojit Sur, Kathleen Gabrielson, Shibin Zhou, Bert Vogelstein, Kenneth W. Kinzler

## Abstract

Chimeric antigen receptors (CARs) can induce T cells to kill cancer cells but also to kill normal cells that express the same antigens^1^. Designing CARs to recognize combinations of antigens, via Boolean logic, can simultaneously expand the scope of targetable antigens and make CAR T cells more specific to cancer^2^. For example, one antigen may be expressed on cancer cells and normal bone marrow cells, while a second antigen may be present on the same cancer cells but only in normal lungs. If recognition of both antigens is required for T cell activation, only the cancer cells will be killed. Creating such AND-gated CAR T cells has been challenging given the need to engineer non-natural signaling mechanisms that integrate two ligand binding events into a single T cell activation stimulus^3–6^. Here, we design a fundamentally new AND-gated receptor called ***M***ulti-***AN***tigen ***T***riggered ***I***mmune ***S***ynapse (MANTIS), which leverages differences in extracellular receptor dimensions to regulate CAR signaling. MANTIS initially prevents CAR activity by steric blocking with a bulky extracellular domain. Upon engagement of the first antigen, MANTIS sheds this blocking domain, releasing a free CAR that can bind a second antigen and activate the T cell in an AND-gated manner. This work demonstrates how differences in extracellular receptor size can be leveraged to spatially regulate intracellular signaling pathways in response to antigen patterns, paving the way for new applications in synthetic biology and cell engineering.

**One sentence summary:** Differences in extracellular size can be leveraged to regulate CAR T cell activity for precise recognition of antigen patterns.

## Main

CAR T cell therapy has produced long-term remissions for patients suffering from B cell cancers^7^. The potential of CAR T cell therapy to treat other cancers is limited by the absence of cancer specific antigens that can be safely targeted^1^. Most candidate antigens are also expressed at lower levels on critical normal tissues, resulting in CAR T cell mediated on-target, off-tumor toxicity^8–11^. Engineering CAR T cells to target combinations of antigens, rather than single antigens, could substantially expand the strategies available to specifically target cancer cells^2^. However, designing T cells to combinatorially recognize antigen patterns has been challenging.

Previously, we and others have investigated the use of NOT-gated logic in CAR T cells to prevent normal cell toxicity^12–15^. In this approach, simultaneous co-engagement of a CAR and an inhibitory CAR (iCAR) on normal cells can prevent T cell activation and killing. Similarly, AND-gated CAR T cells that are activated upon simultaneous engagement of two target antigens have been described^3,4,6^. A potential problem with these innovative approaches is that they may require target antigens to be co-localized at sufficient density and have geometries on the cell surface that enable co-localization of two receptor components^16^.

Rather than receptor co-engagement, CAR logic can also be created via serial engagement, where receptors bind one after the other rather than simultaneously. In one ingenious approach, ligation of a constitutively expressed synthetic NOTCH (SynNOTCH) receptor^17^ to one antigen yields conditional expression of a CAR directed to the second antigen^5,18–20^. Once expressed, the CAR can function independent of the SynNOTCH receptor, meaning that this approach does not require the target antigens to co-localize, and thus is less sensitive to antigen geometries via its IF-THEN operation. However, for some targets, the relatively slow kinetics of transcriptional activation and deactivation render this circuit unable to distinguish between cancer cells expressing both target antigens and two separate normal cells in proximity each bearing a single target^4,21^. Rather than relying on transcriptional regulation, we sought to develop a new AND-gated CAR with improved specificity that functioned via the serial engagement of receptors on the cell surface.

We were inspired by the spatial mechanisms of T cell signaling on the cell surface. T cell signaling is initiated when intracellular membrane-proximal kinases phosphorylate the T cell receptor (TCR) upon ligand binding^22^. To prevent activation in the absence of a ligand, T cells express large quantities of the cell surface phosphatase called CD45, which counteracts TCR phosphorylation^23–25^. CD45 has a bulky extracellular domain (ECD) ∼50 nm in length^26,27^ which upon TCR-ligand binding is excluded from the immune synapse (∼15 nm in length^28,29^) due to size^30–32^. This ensures the synapse is free of inhibitory phosphatases, allowing intracellular kinases to phosphorylate the TCR and initiate T cell activation. This kinetic segregation model of T cell activation can explain how the phosphatase activity of CD45 is regulated by its bulky ECD^33,34^.

We asked whether the ECD of CD45 could be re-wired to control kinase signaling and T cell activation directly based on size rather than phosphatase activity. Coupling steric blocking and antigen dependent proteolysis, we were able to develop a new type of AND-gated CAR that functions via the serial engagement of receptors on the cell surface.

### The CD45 ECD can block CAR activity

CAR and TCR signaling are sensitive to the dimensions of the immune synapse. Elongating receptor-ligand interactions results in weak TCR^35^ and CAR^36^ signaling due to poor CD45 exclusion. Recognizing the spatial nature of T cell signaling (Extended Data Fig. 1a), we hypothesized that CAR signaling could be blocked artificially by coupling to the large ECD of CD45. To test this hypothesis, we attempted to inhibit the activity a CD19-41BBz CAR which can potently kill CD19+ NALM6 target cells^37^. We coupled the ECD of CD45 to the CD19 CAR in two different ways. In the first approach, the CD19 CAR and the CD45 ECD were bound together intracellularly, but not covalently, through a leucine zipper pair^3,38,39^ (Fig. 1a, 45ECD-CAR-1). As this interaction was non-covalent, we were mindful that the dynamic on-off kinetics of the leucine zipper pair could prevent efficient CAR inhibition. In the second approach, the CD19 CAR and the CD45 ECD were joined through a covalent polypeptide linkage (Fig. 1a, 45ECD-CAR-2). We took advantage of the observation that a type II transmembrane protein (intracellular N terminus) can be connected in series with a type I transmembrane protein (intracellular C terminus) and still retain its function^40,41^. Using the type II transmembrane domain from TM11D^40^, we created a single polypeptide where the CD45 ECD in type I orientation was directly coupled to a CD19-41BBz CAR in type II orientation (Fig. 1a, 45ECD-CAR-2). In this design, the CAR signaling domains are inverted relative to their native type I transmembrane orientations. As controls, we created identical CARs without a CD45 ECD (Fig. 1a, CAR-1 & CAR-2).

**Fig. 1.**
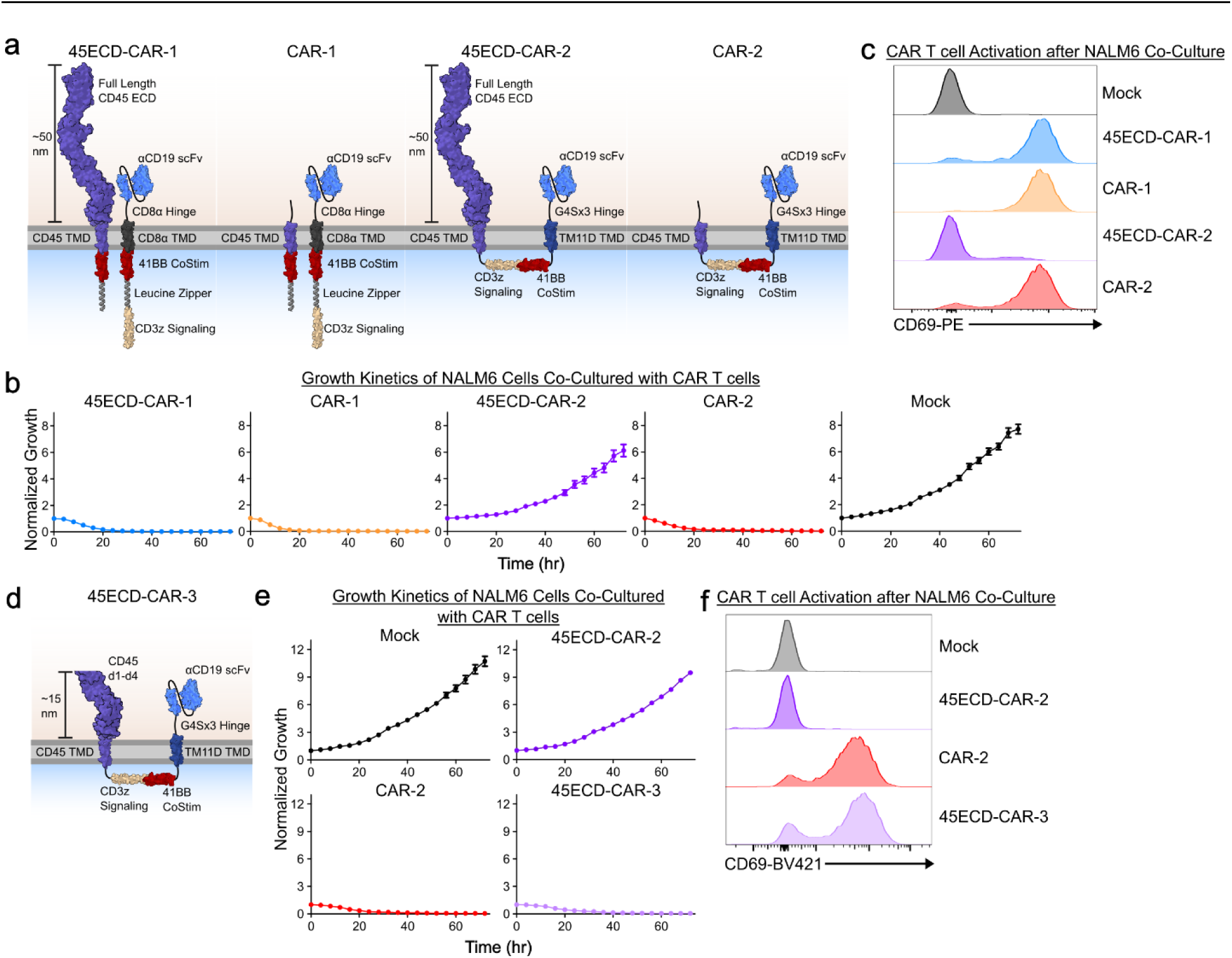
Inhibiting CAR activity with CD45 ECD. **(a)** Design of CD19bbz CARs coupled to full length CD45 ECD in different formats along with control CAR designs. **(b)** 100,000 45ECD-CAR variants or control CAR T cells were co-cultured with 20,000 NALM6 Parental GFP-luciferase cells (5:1 E:T) for 72 hours and tracked by time-lapse imaging. Normalized Growth represents total GFP integrated object intensity normalized by the integrated intensity at the zero hr timepoint. **(c)** 50,000 45ECD-CAR variants and controls were co-cultured with 100,000 NALM6 parental cells (1:2 E:T) for 24 hours and T cell activation was assessed via CD69 expression. **(d)** Design of 45ECD-CAR-3, with a truncated 45ECD containing only d1 to d4. **(e)** Target cell growth kinetics of 45ECD-CAR-3 as tracked via time-lapse imaging as in **b**. **(f)** CD69 activation of 45ECD-CAR-3 and controls as in **c**. Data in **b, c, e,** and **f** are representative of n = 3 technical replicates. The mean +/− standard deviation of technical replicates are plotted in **b** and **e**.

The genes encoding these proteins were introduced into the *TRAC* locus of primary human T cells using CRISPR-mediated homology directed repair^42^ (Extended Data Fig. 1b) and the CARs expressed successfully (Extended Data Fig. 1c). The resultant CAR T cells were then co-cultured in excess with NALM6 target cells expressing GFP-luciferase and monitored by time-lapse imaging. Both the CAR-1 and CAR-2 control T cells, as well as the 45ECD-CAR-1 T cells, potently killed the NALM6 targets cells, but 45ECD-CAR-2 T cells did not affect NALM6 growth (Fig. 1b). This result was confirmed by independent assays at a variety of effector-to-target cell (E:T) ratios (Extended Data Fig. 1d). To determine whether the inhibitory effect was due to interference with CAR T cell activation, we assessed CD69 expression (a protein associated with T cell activation) in the same cells. All CAR T cells except 45ECD-CAR-2 cells were found to be activated when incubated with NALM6 cells (Fig. 1c).

To further investigate the mechanism of CAR T cell inhibition by CD45 ECD, we generated 45ECD-CAR-2 T cells with endogenous CD45 knocked-out (Extended Data Fig. 1e). Knock-out (KO) of endogenous CD45 did not augment the activation status of 45ECD-CAR-2 expressing T cells upon co-culture with NALM6 cells (Extended Data Fig. 1f), indicating that the inhibitory effect of CD45 ECD did not depend on retention of endogenous CD45 phosphatase. This suggested that the inhibitory effect of CD45 ECD could be due to steric blocking rather than phosphatase inhibition. To further test this idea, we truncated CD45 ECD so that it included only d1 to d4, the rigid membrane proximal portion of CD45 ECD that is only ∼15 nm in length^27^ (Fig. 1d, 45ECD-CAR-3). This is approximately the same length as the CD19 CAR immune synapse (∼10-12 nm^36,43^) and thus we expected no inhibitory effect on CAR signaling. We first verified that 45ECD-CAR-2 and 45ECD-CAR-3 were expressed at similar levels (Extended Data Fig. 1g). When co-cultured with NALM6 cells, the truncated CD45 ECD domain in 45ECD-CAR-3 T cells failed to inhibit CAR activity as observed by time-lapse imaging (Fig. 1e), a luminescence assay (Extended Data Fig. 1h), and via CD69 expression (Fig. 1f). Collectively, these data suggested that full length CD45 ECD could inhibit CAR signaling in a size dependent manner.

### Antigen dependent CD45 ECD proteolysis enables AND-gating

To make a functional AND-gate, we envisioned that CD45 ECD would be proteolyzed in an antigen dependent manner to unleash a “free” CAR that could activate the T cell in response to a second antigen, creating an AND-gate (Fig. 2a). Native CD45 can be proteolyzed in some activated monocytes and granulocytes, but may not be a substrate of proteolysis in T cells^44^. However, ECDs even with non-canonical or unknown protease sites can be designed to undergo efficient antigen-dependent proteolysis when engineered into synthetic intramembrane proteolysis receptors (SNIPRs)^45^. We therefore attempted to convert the CD45 blocking component of 45ECD-CAR-2 to function as a SNIPR that could proteolyze upon antigen engagement. First, we appended a single-chain variable fragment (scFv) to the N-terminus of 45ECD-CAR-2. Then, we replaced the transmembrane (TMD) and intracellular juxtamembrane (JMD) domains of CD45 with those of NOTCH1 and NOTCH2 respectively based on a previously optimized SNIPR design^45^. We named this design: MANTIS (for ***M***ulti-***AN***tigen ***T***riggered ***I***mmune ***S***ynapse). As NALM6 cells express high levels of HLA-A2 (A2), we designed our first MANTIS construct, MANTIS-1, to target cells that express A2 (via the CD45 arm) *AND* CD19 (via the CAR arm) (Fig. 2b, c). Analogous single antigen targeted CAR T cells were used as controls (Fig. 2b, 19CAR & A2CAR). At low E:T ratios, MANTIS-1 T cells preferentially killed the parental NALM6 target cells at similar levels to control CAR T cells (Fig. 2d, Extended Data Fig. 2a,b). Remarkably, at all tested E:T ratios, MANTIS-1 did not kill the CD19 KO (A2+) target cells (Fig. 2d, Extended Data Fig. 2b) meaning MANTIS-1 was not activated by independent binding of A2 via the CD45 arm of the receptor. However, with higher E:T ratios, MANTIS-1 had increasing toxicity to the A2 KO (CD19+) target cells (Fig. 2d, Extended Data Fig. 2b). This suggested that the non-specific activity of MANTIS-1 could be due to CD45 ECD proteolysis independent of A2 binding, releasing free CARs that could activate the T cell upon binding CD19 alone.

**Fig. 2.**
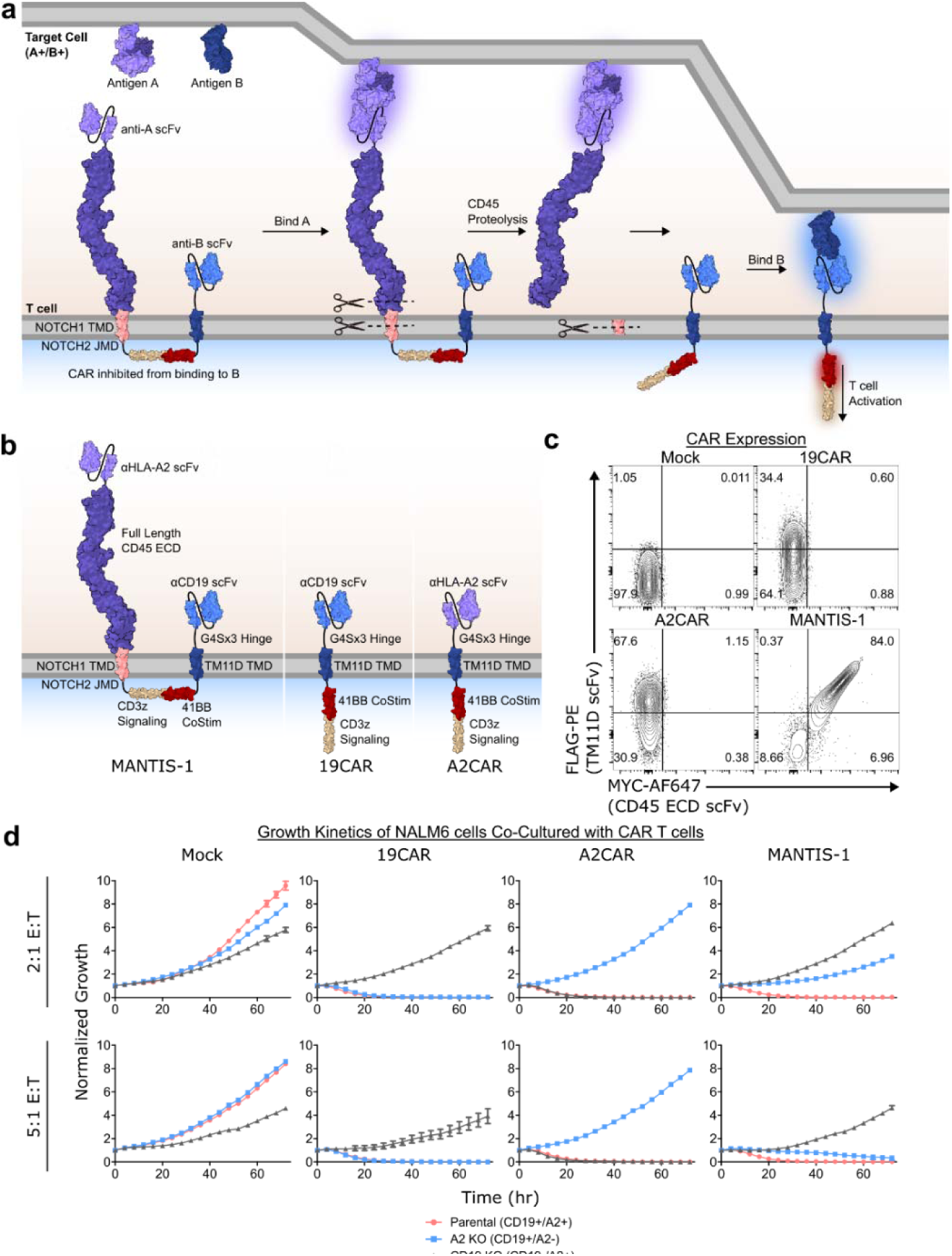
CD45-Proteolysis Creates AND-gated CAR T cells. **(a)** Scheme for MANTIS design in which a CAR is inhibited from binding to its antigen by covalent attachment to the bulky CD45 ECD. MANTIS is designed with an scFv placed N-terminal to the CD45 ECD domain, which upon ligand binding induces CD45 ECD proteolysis. This releases the CAR domain that can subsequently bind to a second antigen and activate the T cell. **(b)** Design of MANTIS-1, which was targeted to cells expressing A2 and CD19. MANTIS-1 was designed with a SNIPR core^45^, consisting of a NOTCH1 TMD and NOTCH2 JMD. CAR controls in type II transmembrane orientation are also depicted. **(c)** Expression of MANTIS-1 and CAR controls as identified by staining for MYC and FLAG tags placed at the N- and C-termini. **(d)** MANTIS-1 and controls were co-cultured with 20,000 NALM6 Parental (CD19+/A2+), A2 KO (CD19+), and CD19 KO (A2+) target cells for 72 hours at different E:T ratios and target cell growth was visualized via time-lapse imaging. Normalized Growth represents total GFP integrated object intensity normalized by the integrated intensity at the zero hr timepoint. Data in **d** represents n = 2 experiments with 3 technical replicates for the 2:1 E:T ratio. For the 5:1 E:T ratio, data represents n = 3 technical replicates. The mean +/− the standard deviation of all technical replicates is plotted.

Using a panel of SNIPR-reporter constructs which induce GFP expression upon antigen-exposure^45^ (Extended Data Fig. 3a), we indeed found that CD45 ECD-NOTCH chimeras were proteolyzed whether or not they were exposed to antigen (Extended Data Fig. 3b-g). This constitutive proteolysis could potentiate the observed non-specific activity by MANTIS-1. Controls for this experiment were provided by SNIPR-3, which contained a CD45 TMD/JMD and did not induce GFP-expression at baseline or upon exposure to antigen (SNIPR-3, Extended Data Fig. 3b-d).

To overcome non-specific activity, we hypothesized that engineering antigen-dependence in a SNIPR-GFP reporter format would yield more specific MANTIS designs. We found that the addition of NOTCH negative regulatory regions (NRRs)^46–48^ to SNIPRs with a CD45 ECD yielded ligand-dependent GFP expression in a NOTCH-like manner (Extended Data Fig. 3h-l). However, NRR addition into MANTIS-1 did not resolve non-specific activity to the A2 KO (CD19+) target cells, even with several modifications including: addition of a protective RAM7 sequence^49^, deletion of constitutive proteolysis sites^50,51^, or incorporation of other NRR variants^52,53^ (MANTIS 2-11, Extended Data Fig. 4).

### Designing a MANTIS without any NOTCH components

NOTCH proteolysis can be enhanced by T cell activation^45,54^, and our failures to improve MANTIS specificity by modifying NOTCH domains (Extended Data Fig. 4) prompted us to eliminate all NOTCH components from MANTIS-1. We started by replacing the TMD and JMD of MANTIS-1 with those of CD45, creating MANTIS-12 (Fig. 3a,b). We expected MANTIS-12 to have reduced cytolytic activity because a SNIPR reporter construct (SNIPR-3) that contained a CD45 TMD and JMD (instead of NOTCH1 TMD and NOTCH2 JMD) was inactive (Extended Data Fig. 3b-d). To our surprise, MANTIS-12 elicited highly specific AND-gated activity as measured by multiple metrics of cell killing and T cell activation even when incubated in vast excess to target cells (Fig. 3c-f).

**Fig. 3.**
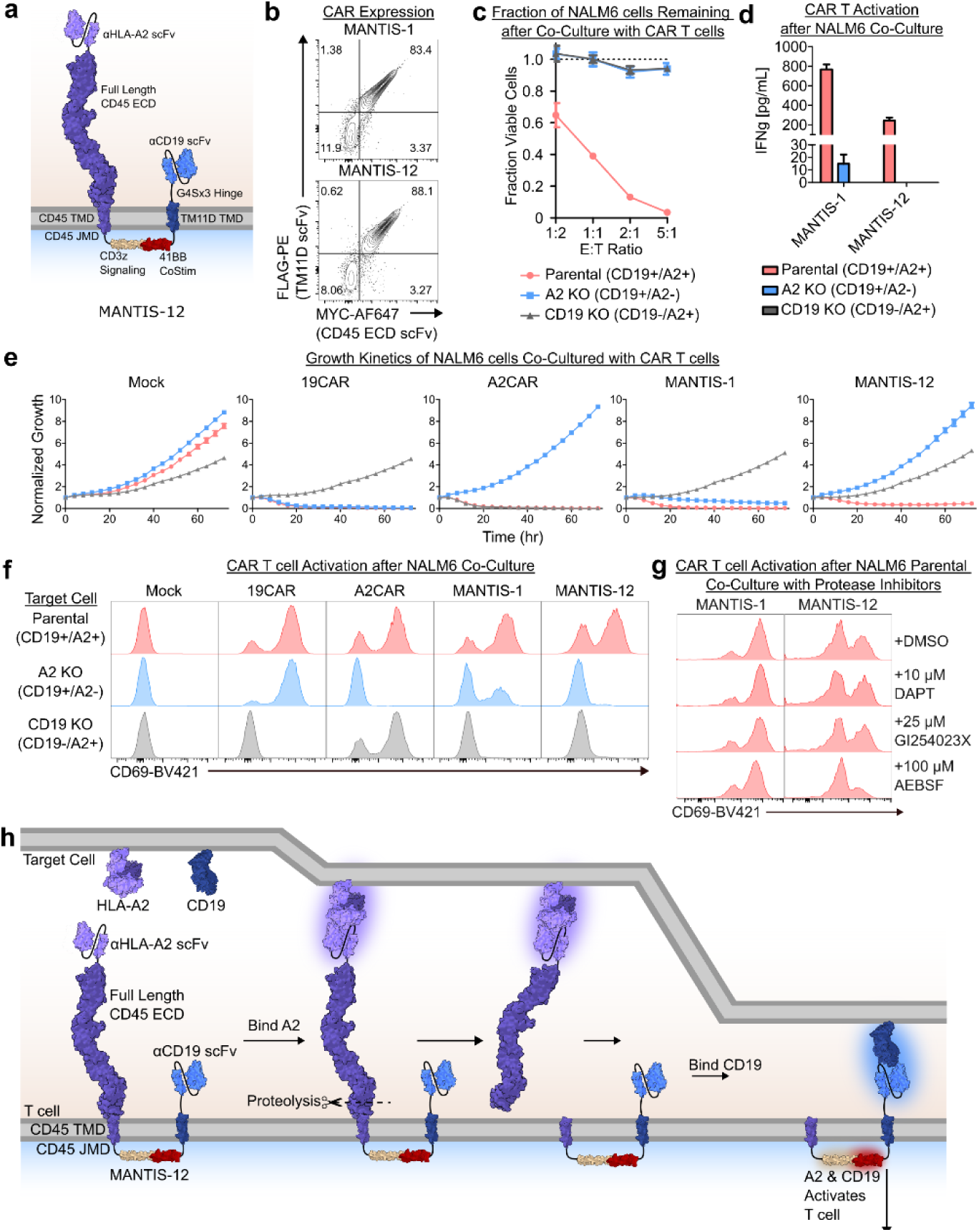
MANTIS specificity can be improved by incorporating a native CD45 transmembrane anchor. **(a)** MANTIS-12 design, which was created by replacing the NOTCH1 TMD and NOTCH2 JMD in MANTIS-1 with CD45 TMD and JMD, respectively. **(b)** Expression of MANTIS-12 and controls as identified by staining for MYC and FLAG tags placed at the N- and C-termini. **(c)** 5,000-50,000 MANTIS-12 cells and controls were co-cultured with 10,000 of each NALM6 GFP-luciferase isogenic lines in individual wells. After 48 hrs, the fraction of viable cells was measured by a SteadyGlo luminescence assay. The fraction of viable cells compared to the Mock condition is depicted. **(d)** IFNg secretion by MANTIS-12 and MANTIS-1 cells after co-culture at a 1:2 E:T ratio with 100,000 NALM6 isogenic cell lines in individual wells. Negative values were converted to zero. **(e)** 20,000 MANTIS-12 and control T cells were co-cultured with NALM6 target cells at a 5:1 E:T ratio and target cell growth was measured by time-lapse imaging. Normalized Growth represents total GFP integrated object intensity normalized by the integrated intensity at the zero hr timepoint. **(f)** CD69 expression on MANTIS cells and CAR controls when co-cultured with 100,000 NALM6 isogenic cell lines in a 1:2 E:T ratio for 24 hours. **(g)** 20,000 MANTIS-1 and MANTIS-12 cells were co-cultured with 20,000 NALM6 Parental (CD19+/A2+) cells in a round-bottom 96-well plate in the presence of a gamma secretase inhibitor (DAPT), ADAM inhibitor (GI254023X), and serine protease inhibitor (AEBSF) for 4 hours. After culture, T cell activation was assessed via CD69 expression. **(h)** Mechanism of activation by MANTIS-12 based on observations that CD45 ECD can be shed as shown in **g** and CD45 TMD/JMD is not proteolyzed. Data in **c**, **e,** and **g** are representative of n = 3 technical replicates. Data in **d** and **f** are representative of n = 2 experiments with 3 technical replicates. The mean +/− standard deviation of replicates are plotted in **d** and **e**.

To determine whether MANTIS-12 produced logic-gated activity via the proteolysis of the CD45 ECD, we tested the impacts of various protease inhibitors on MANTIS-12 activation when in co-culture with NALM6 parental cells (Fig. 3g). Only AEBSF, a serine protease inhibitor, reduced the activation of MANTIS-12 T cells (Fig. 3g). This was in line with previous observations that AEBSF prevented CD45 proteolysis in monocytes^44^. Note that MANTIS-1 T cells were not inhibited by AEBSF or two other protease inhibitors (the ADAM inhibitor: GI254023X and the gamma secretase inhibitor: DAPT). This was consistent with our postulated mechanism that the non-specific activity of MANTIS-1 was driven by baseline antigen-independent proteolysis of CD45 ECD. We further verified the ability of CD45 ECD to be shed from MYC-tagged MANTIS-12 expressed in Jurkat cells in response to anti-MYC bead stimulation (Extended Data Fig. 5). This suggested that CD45 ECD shedding by MANTIS-12 depended only on binding-triggered stimulation of the CD45 ECD arm. Collectively, these results showed that MANTIS-12 produces AND-gated activity via the antigen-dependent shedding of CD45 ECD (Fig. 3h).

### MANTIS Retains Specificity in Mixed Cell Models

MANTIS-12 T cells could accurately distinguish double-target from single-target cells in isolated co-cultures (Fig. 3c-f). We sought to understand whether MANTIS-12 retained specificity in mixed culture systems. First, we tested whether MANTIS-12 could prime on one cell expressing a single target antigen and then kill a nearby cell expressing the other target antigen. This was an undesirable behavior a previously described IF-THEN receptor circuit^4,21^. We co-cultured MANTIS-12 T cells with the NALM6 CD19 KO and A2 KO target cells mixed in the same well at equal ratios (Fig. 4a). As hoped, MANTIS-12 T cells were not activated by the expression of the two antigens on separate target cells and did not induce any cell killing unlike MANTIS-1 and control CAR T cells, which mediated on-target and bystander killing (Fig. 4b-c).

**Fig. 4.**
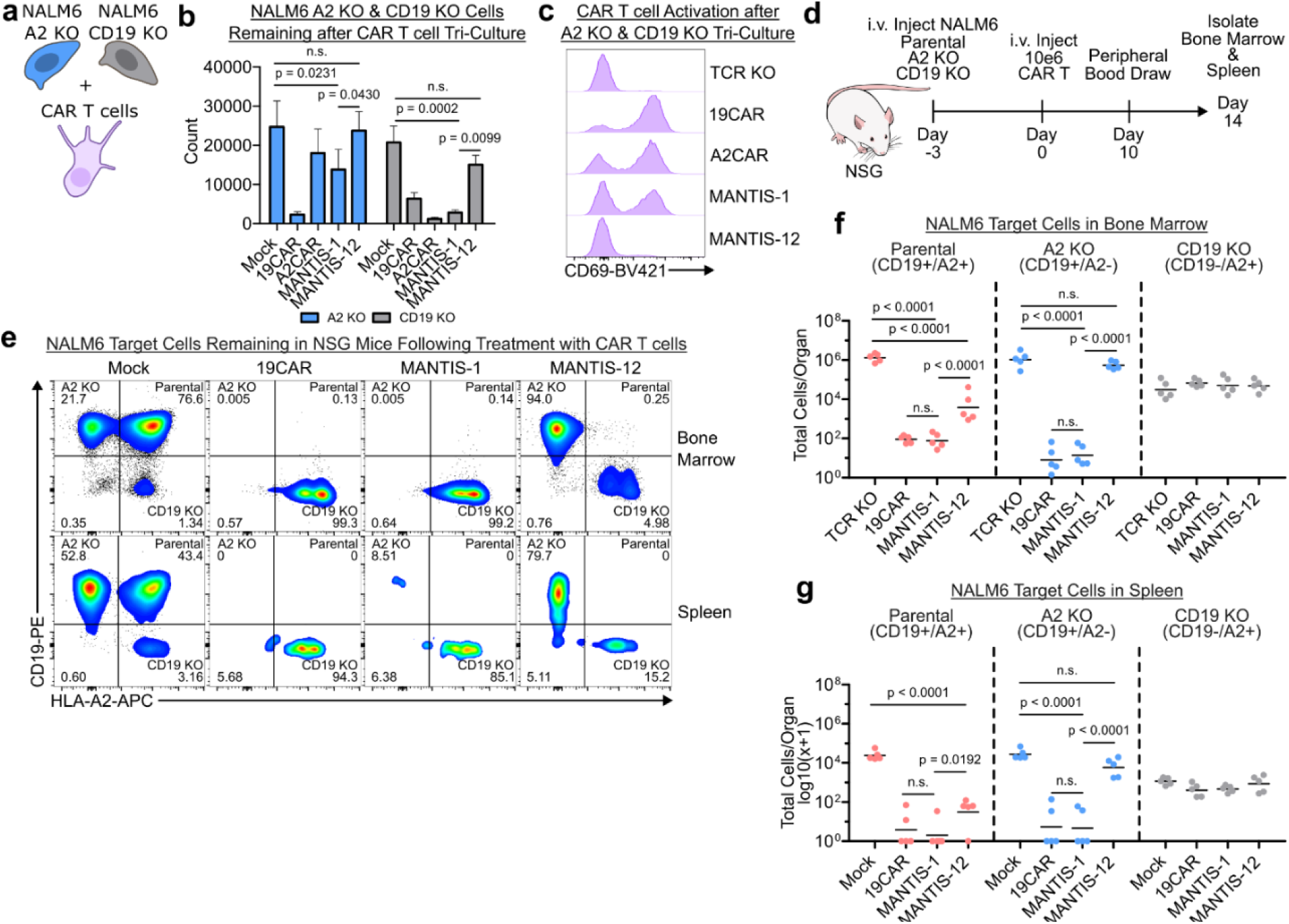
MANTIS exhibits specificity in mixed culture models. **(a)** 25,000 MANTIS and control CAR T cells were co-cultured with single target NALM6 A2 KO (CD19+) and CD19 KO (A2+) cells in the same well in a 1:1:1 ratio. **(b)** After 24 hours in co-culture, residual target cells were counted using flow cytometry. p values were calculated with a two-way ANOVA with Tukey’s multiple comparisons test. **(c)** CAR T cell activation was assessed by CD69 flow staining. (n = 2 experiments for **b** and **c** with three technical replicates). The mean and standard deviation of technical replicates are plotted in **c**. **(d)** Mixed target cells *in vivo* model. 100,000 NALM6 Parental, 100,000 NALM6 A2 KO, and 400,000 CD19 KO cells were intravenously injected in NSG mice 3 days prior to treatment with 10 million CAR or MANTIS T cells. Ten days after CAR administration, T cell engraftment was assessed via flow cytometry and 14 days after treatment target cells were assessed via flow cytometry. **(e)** Representative flow plots showing target cells in the bone marrow and spleen of NSG mice after 14 days of treatment with CAR or MANTIS cells. Count of target cells isolated from **(f)** bone marrow and **(g)** spleens of NSG mice treated with CAR and MANTIS cells (n=5 mice per group, every data represents one animal; p values calculated by a two-way ANOVA with Tukey’s multiple comparisons test of log-transformed values). Bars represent the mean of each replicate.

To evaluate if MANTIS-12 T cells could reliably distinguish double-target from single-target cells *in vivo*, we injected all three NALM6 target cells intravenously into immunocompromised mice and treated them with a high dose of CAR T cells (Fig. 4d, Extended Data Fig. 6a-c). CAR T cell engraftment was verified via flow cytometry (Extended Data Fig. 6d-f). As MANTIS T cell treatment should not be curative in this model because of the inclusion of non-targeted single antigen cells, we sacrificed the mice 14 days after treatment to identify engrafted tumor cells in the bone marrow and spleen. MANTIS-1 and the CD19 CAR T cells showed similar activity, in that they depleted CD19 expressing cells in both bone marrow and spleens (Fig. 4e-g). In contrast, MANTIS-12 T cells specifically depleted only the double-target parental NALM6 cells and spared all single target cells (Fig. 4e-g).

### Optimizing MANTIS for the Treatment of Solid Tumors

CAR T cells targeted to several solid tumor antigens have yielded severe on-target off-tumor toxicities in humans^8–11^. A classic example is the case of a patient treated with anti-HER2 CAR T cells who suffered lethal pulmonary toxicity^9^. Similarly, severe pneumonitis was observed in mice treated with CAR T cells directed by a cross-reactive anti-HER2 DARPin (H10-2-G3)^55,56^. Logic-gated CAR T cells have the potential to avoid this toxicity^56^, and we sought to test whether MANTIS T cells could overcome on-target off-tumor toxicity in this pre-clinical model.

We picked the ovarian adenocarcinoma cell line, SKOV3, as a model target cell because of its high expression of HER2 and susceptibility to anti-HER2 CAR T cells^57^. SKOV3 cells also express high amounts of EGFR, thus we chose EGFR *AND* HER2 as a model antigen pair. To address receptor specificity, we engineered EGFR KO and HER2 partial KO (pKO) variants of SKOV3 (Extended Data Fig. 7a). A complete HER2 KO could not be generated possibly due to the high dependency of SKOV3 on HER2 for survival^58,59^.

In prior reports, lung toxicity in mice was better potentiated by DARPin CARs with CD28 co-stimulation^55^. Thus, we modified MANTIS-12 to contain CD28 co-stimulation and re-targeted the receptor to EGFR (via the CD45 arm) *AND* HER2 (via the CAR arm), creating MANTIS-13 (Extended Data Fig. 7b). All constructs and control CARs successfully expressed in primary T cells (Extended Data Fig. 7b-c). In co-culture, MANTIS-13 more potently killed the EGFR+/HER2+ SKOV3 parental cells over single target cells at a 5:1 E:T ratio (Extended Data Fig. 7d). However, SKOV3 parental killing was not complete, and MANTIS-13 also had moderate reactivity to the HER2 pKO (EGFR+) target cells. To orthogonally evaluate this non-specific activity, we overexpressed EGFR and HER2 in Jurkat cells which naturally do not express these proteins (Extended Data Fig. 7e, Methods). Co-culture with Jurkat target cells revealed that MANTIS-13 T cells could kill EGFR overexpressing Jurkat cells (Extended Fig. 7f-g). Collectively, this suggested that MANTIS-13 could be stimulated by independent binding of the anti-EGFR scFv on the CD45 ECD arm. This was distinct from the CAR-mediated non-specific activity we observed with MANTIS-1, and thus, we sought to further optimize MANTIS-13.

We first enhanced CAR potency by adding a MyD88-CD40 (MC) co-stimulation module^60,61^ given our prior experience^62^. Alongside MC, we added an additional CD3z signaling domain in the type I orientation by using a pair of anti-parallel leucine zippers^63^ (Extended Data Fig. 8a) since the non-native type II orientation of the CD3z domain in the CAR portion of the original MANTIS could potentially impair efficient signaling. Adding this accessory signaling module slightly decreased CAR expression (HER2CAR-MC Extended Data Fig. 8b), but substantially improved CAR potency (Extended Data Fig. 8c-d).

Regarding specificity, previous reports have suggested that elongated immune synapses could signal albeit at a reduced efficiently, given a high enough affinity receptor-ligand interaction^64,65^. This suggested that MANTIS receptors engineered with high affinity N-terminal binders may result in non-specific T cell activation. MANTIS-12 retained high specificity (Fig. 3-4) and the use of a moderate affinity N-terminal scFv (BB7.2, 162 nM^66^) may support this hypothesis. We then created MANTIS-MC receptors and sought to reduce the affinity of the anti-EGFR scFv on the CD45 arm using variants derived from the C10 antibody clone^67^ (MANTIS 14-17, Extended Data Fig. 8e). In these experiments, we used the human-specific 4D5 scFv as the anti-HER2 binder. We co-cultured the new MANTIS receptors (Extended Data Fig. 8e, f) with the engineered Jurkat target cells because they were more sensitive to killing. MANTIS variants with lower affinity N-terminal binders were more specific, and the lowest affinity variant (MANTIS-17) had near perfect specificity (Extended Data Fig. 8g-h).

### MANTIS Performance in Models of On-Target Off-Tumor Toxicity

To study safety and efficacy, we then changed 4D5 in MANTIS-14 and 17 to the H10-2-G3 DARPin creating MANTIS-18 and 19 (Fig. 5a-b). As a benchmark, we also modified a previously described AND-gated receptor, LINK CAR^4^, to target EGFR and HER2 (Fig. 5a-b). Both MANTIS T cell variants could completely clear SKOV3 parental cells *in vitro*, and the lower affinity MANTIS-19 variant retained perfect AND-gated specificity (Fig. 5c-d) as previously observed in our Jurkat optimization (Extended Data Fig. 8e-f). In contrast, LINK CAR could not mediate any SKOV3 lysis at a 5:1 E:T ratio (Fig. 5c-d), possibly due to the lack of additional co-stimulation required for solid tumor activity^68^. LINK CAR could mediate target lysis of Jurkat cells overexpressing EGFR and HER2 (Extended Data Fig. 9a-c), but also had some leaky activity based on the LAT component of the receptor as previously reported^4^ (Extended Data Fig. 9b-c).

**Fig. 5.**
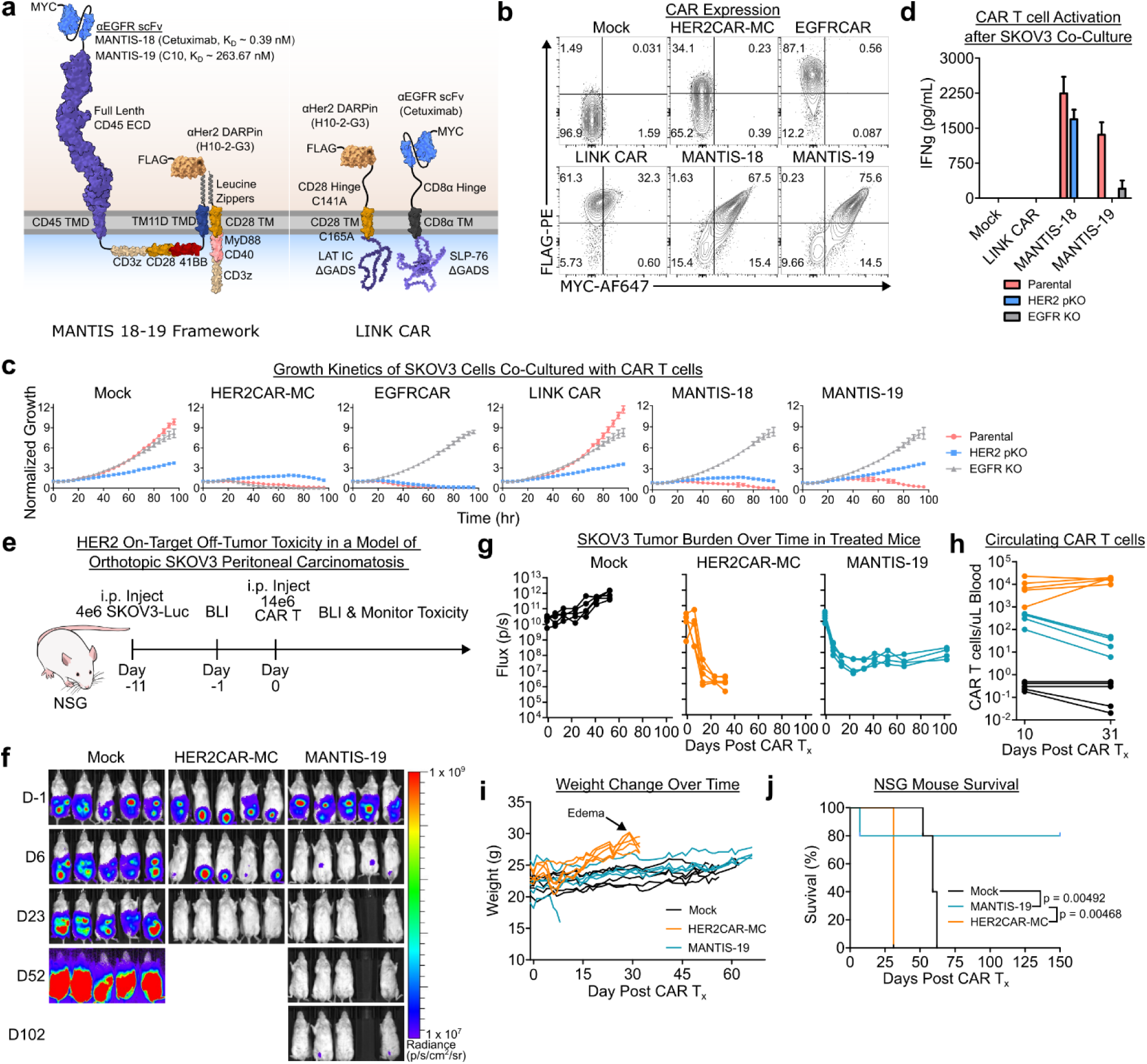
MANTIS performance in a model of HER2 on-target off-tumor toxicity. **(a)** Design of MANTIS-18, MANTIS-19, and LINK CAR T cells. **(b)** Expression of CARs as identified by cell surface staining for MYC and FLAG tag. EGFRCAR and HER2CAR-MC encode only the FLAG tag. **(c)** CAR T cells in **b** were co-cultured with 2,000 SKOV3 cell lines at a 5:1 E:T ratio and target cell growth was monitored with time-lapse imaging. Normalized Growth represents total GFP integrated object intensity normalized by the integrated intensity at the zero hr timepoint. **(d)** IFNg secretion by CAR T cells depicted in **a** from the co-culture depicted in **c** at the 72-hour timepoint. Negative values converted to zero. **(e)** Experimental timeline for the NSG mouse experiment to study HER2 on-target off-tumor toxicity with SKOV3 xenografts implanted in the peritoneum. Mice were treated i.p. with a high dose of Mock, HER2CAR-MC, or MANTIS-19 cells normalized to 90% NGFR purity. **(f)** Anti-tumor response to CAR T cell therapy as visualized by bioluminescence imaging. **(g)** Quantified tumor burden in NSG mice over time. **(h)** Flow cytometric quantification of circulating CAR T cells in blood. **(i)** Weight change and **(j)** survival of NSG mice after various treatments over time. p values calculated using the Fleming-Harrington weighted log-rank test. Data in **c** and **d** are representative of n = 3 technical replicates. The mean and standard deviation of all replicates is plotted. Data in **f-i** representative of n = 5 mice per group. Data for MANTIS-19 in **h** represent only of n = 4 mice. BLI = bioluminescence imaging. T_x_ = treatment.

Seeing the high potency and specificity of MANTIS-19 *in vitro*, we tested this construct with *in vivo* models of HER2 on-target off-tumor toxicity^55^ (Fig. 5e). First, we orthotopically implanted luciferase expressing SKOV3 parental cells (SKOV3-luc) by intraperitoneal (i.p.) injection in immunodeficient mice and verified tumor engraftment ten days after injection. After randomization the following day, mice were treated with an i.p. injection of CAR T cells (Extended Data Fig. 10a) based on prior reports that local administration of CAR T cells is more effective in models of peritoneal carcinomatosis^69–72^. HER2CAR-MC and MANTIS-19 T cells rapidly induced tumor regression (Fig. 5f-g). CAR T cells, for all conditions, entered systemic circulation by Day 10 post i.p. injection, but only HER2CAR-MC T cells were expanding by day 31 post injection (Fig. 5h). After 3 weeks of treatment, HER2CAR-MC treated mice developed progressively worsening edema involving hind legs, tail, and face resulting in weight gain and 5/5 mice were euthanized (Fig. 5i). On histopathological examination, all HER2CAR-MC mice demonstrated severe lymphocytic infiltration in the lungs (Extended Data Fig. 10b) indicative on-target off-tumor toxicity. One of the MANTIS-19 mice died early on day 7 due to liver necrosis (Extended Data Fig. 10c) and demonstrated mild lymphocytic infiltration in the lungs (Extended Data Fig. 10b). This phenotype may be due to the MC module incorporated into MANTIS-19, as a similar MC module has been shown to induce cytokine release syndrome (CRS) in other models^60^. Remarkably, 4/5 MANTIS-19 treated mice demonstrated strong anti-tumor responses and extended survival beyond 150 days post-treatment without any signs of toxicity (Fig. 5i,j).

Finally, to enable comparison to LINK CAR, we tested safety and efficacy in a Jurkat model of HER2 on-target off-tumor toxicity. Two days post cancer cell injection, NSG mice were treated with an i.v. injection of 10 million CAR T cells (Extended Data Fig. 11a-b). Mock treated mice rapidly succumbed to neoplastic disease with a median survival of 33 days (Extended Data Fig. 11c-f). HER2CAR-MC, MANTIS-19, and LINK CAR all induced an immediate anti-tumor response, but cancer recurred in 2/5 LINK CAR mice (Extended Data Fig. 11c-d). In this experiment, HER2CAR-MC mice developed worsening edema by day 38 following treatment. These mice eventually experienced labored breathing and weight loss due to HER2 on-target off-tumor toxicity and were euthanized. The MANTIS-19 treated mice lost weight immediately after treatment, but these mice recovered and showed a durable anti-cancer effect (See Methods, Extended Data Fig. 11c-f). Altogether, these results showed that MANTIS could minimize HER2 on-target off-tumor toxicity without compromising anti-tumor efficacy across multiple models.

## Discussion

Combinatorial antigen targeting is expected be critical for CAR T cell therapy to be impactful beyond B cell malignancies^2^. AND-gated CARs enable T cells to precisely kill target cells based on the high expression of two target antigens, reducing on-target off-tumor toxicity. However, the need to convert dual-antigen binding into a single activation signal is challenging and there are only a small number of studies describing AND-gated CAR T cells^3–6^. We created an entirely new type of AND-gated CAR named MANTIS which was inspired by the kinetic segregation model of T cell activation^33^. Some groups have leveraged CD45 to modulate immune responses^73–75^. MANTIS rewires the ECD of CD45, which naturally controls intracellular phosphatase activity^34^, to regulate CAR signaling directly. To achieve this, we modified CD45 ECD in such a way that it initially inhibits CAR binding by steric blocking. Binding to the first antigen triggers CD45 ECD shedding, releasing a free CAR that can bind to a second antigen and activate the T cell in a logic-gated manner.

Much like the SynNOTCH circuit^5^, MANTIS operates via the serial engagement of receptors in an IF-THEN manner. One key difference is that MANTIS executes its logic-gated activity on the cell surface, rather than inside the cell via transcription. This overcomes a potential problem with the transcriptional circuit, where the slow kinetics of operation enable SynNOTCH T cells to activate its circuit on one cell, detach from that cell, and then non-specifically kill another cell^4,21^. MANTIS T cells did not demonstrate this behavior in the NALM6 mixed target cell model, though it may potentially be an issue when MANTIS T cells are simultaneously conjugated to multiple target cells. Despite these drawbacks, MANTIS appeared to out-perform a previously described gold-standard AND-gate, LINK CAR^4^, in relevant models of EGFR & HER2 targeting. However, it is worth noting that we needed to titrate the scFv affinity and increase signaling strength on MANTIS to achieve an optimal balance between potency and specificity. Even so, the current MANTIS design is a proof-of-concept and will require further optimization given our observations of initial toxicity possibly resulting from the MC module^60^. Several elements in the MANTIS design can be modified including the bulky extracellular domain, membrane linkages, and signaling domains. This highlights the increased complexity of logic-gated receptors compared to standard CARs, and a similar optimization may also be required to optimize the activity of other AND-gated CARs. Each antigen pair is unique, and some AND-gated receptors may have distinct advantages over others in different situations. Further research is necessary to identify optimal antigen pairs which can be safely and effectively targeted by MANTIS and other logic-gated receptors to treat cancer and other diseases.

In a broader context, MANTIS illustrates how differences in ECD size can be leveraged to control intracellular receptor activity in response to antigen patterns. In the process of engineering MANTIS, we also demonstrated how different CAR modules can be linked covalently by coupling type I and type II membrane proteins. In line with other observations^45^, we further showed how ECD domains without canonical protease sites can be proteolyzed in response to antigen engagement. Although we engineered MANTIS with the intent of creating logic-gated CARs, the MANTIS framework can be easily adapted to regulate the activity of other receptors that rely on antigen binding to control downstream cell signaling, transcription, or other functions in response to antigen patterns. Ultimately, MANTIS not only expands the tool kit of logic-gated CARs which can be used for precisely targeting pathogenic cells but may also serve as a blueprint for engineering basic Boolean operations in biological systems.

## Material & Methods

### CAR Cloning & Construction

CAR and SNIPR amino acids were codon optimized by GeneArt and ordered as GeneArt Strings DNA Fragments. Some fragments were also ordered as gBlocks from IDT DNA. CAR DNA fragments were cloned into a modified pUC19 vector backbone using the NEBuilder HiFi DNA Assembly Cloning Kit (NEB, #E5520S) according to manufacturer protocols. This pUC19 vector encoded 3’ and 5’ homology arms for *TRAC* knock-in, an EF1a promoter, and a 2A-truncated NGFR (tNGFR, AA 1-277) tag for cell sorting. Plasmid clones were sequence verified using either Sanger Sequencing or Whole Plasmid Sequencing (Plasmid-EZ, Genewiz). MANTIS variants were constructed using the full length CD45 extracellular domain (PTPRC RABC isoform, AA26-577) and the type II-oriented membrane anchor from TM11D (AA2-49). For some MANTIS designs, the CD45 TMD (AA578-598) and/or JMD were used (AA599-608). MANTIS 1-12 receptors were constructed with the BB7.2 (anti-A2) and fmc63 (anti-CD19) scFvs. MANTIS 13-19 receptors were constructed with either the H10-2-G3 DARPin or 4D5 scFv for the anti-HER2 binder. ScFvs derived from cetuximab or the C10 clone were used as the anti-EGFR binders. SNIPR and LINK CAR constructs were assembled using previously reported amino acid sequences^4,45^.

### Homology Directed Repair Template (HDRT) Synthesis

Sequence verified plasmids were converted into double stranded DNA homology directed repair templates (HDRTs) via PCR using the NEB Q5 Hot Start High-Fidelity Master Mix (NEB, #M0494X). For each construct, PCR reactions were pooled and purified using Ampure XP beads (Beckman Coulter Life Sciences, #A63880). In brief, 550 μL of PCR product was mixed with 550 μL of Ampure XP beads at room temperature for 10 minutes. DNA bound to beads were collected on a magnet and washed twice with 70% ethanol. HDRTs were eluted in 25 μL nuclease free water at a concentration of 0.5-2 μg/μL, as measured by Nanodrop spectrophotometer (ThermoFisher Scientific).

### T cell Culture & Engineering

Primary human T cells were derived from commercially available leukapheresis products (StemCell Technologies & AllCells). Peripheral blood mononuclear cells (PBMCs) were purified from apheresis products using density gradient centrifugation with either Ficoll Paque Plus (Cytiva, #17144002) or Lymphoprep (StemCell Technologies, #18060). Isolated PBMCs were cryopreserved in 90% FBS, 10% DMSO and kept in liquid nitrogen cold storage until use for engineering purposes.

T cells were engineered in a similar way to previously reported studies^42,62,76^. Two days prior to electroporation, T cells were isolated from frozen PBMCs via immunomagnetic isolation using the EasySep Human T cell Isolation Kit (StemCell Technologies, #17951) and then stimulated at a density of 1.5e6 cells/mL with CD3/CD28 Dynabeads (Gibco, #11131D) in a 1:1 ratio in T cell media (RPMI-1640 supplemented with 10% FBS, 1% P/S, recombinant human IL-2 (100 IU/mL, Proleukin, Prometheus Laboratories), and recombinant human IL-7 (5 ng/mL, Biolegend, #581908)) for 48-56 hours. Following CD3/CD28 stimulation, Dynabeads were removed from activated T cells using a magnet and rested at room temperature. During this time, the electroporation substrate was prepared. Every 1e6 T cells were electroporated with 25 pmol of *TRAC* targeting A.s. Cas12a Ultra (Cpf1, IDT, #10001273) RNP, 25 pmol of *TRBC* targeting RNP, and 500 ng of HDRT. Briefly, *TRAC* guide RNA (GAGTCTCTCAGCTGGTACAC, IDT DNA), *TRBC* guide RNA (GCCCTATCCTGGGTCCACTC, IDT DNA), and Cpf1 electroporation enhancer (IDT, #1076301) were resuspended in IDT Duplex Buffer (IDT, #11-01-03-01) at 100 μM concentration. To prepare 25 pmol of *TRAC* or *TRBC* RNP, 25 pmol of cpf1 enzyme was complexed with 50 pmol of guide RNA and 37.5 pmol of electroporation enhancer for 15 minutes. *TRAC* and *TRBC* targeting RNP master mixes were prepared in separate tubes, and after incubation were mixed in a 1:1 ratio. In experiments with endogenous CD45 KO, an additional 25 pmol of Cas9 RNP was prepared as above and included in the electroporation. Immediately prior to electroporation, T cells were centrifuged at 90 g for 10 minutes and then resuspended in P3 buffer (Lonza, #V4XP-3034) at a density of 50e6 cells/mL. For a typical experiment, 5e6 T cells in 100 μL P3 buffer were mixed with 12.75 μL *TRAC/TRBC* RNP, and 2.5 μg of HDRT diluted in 5 uL in duplex buffer. 110 μL of this mixture was transferred into a 100 uL cuvette and electroporated using pulse code EO-115 on the Lonza 4D Nucleofector X-Unit (Lonza, AAF-1003X). After electroporation, T cells were rested at room temperature for 10 minutes, after which 500 uL of warm RPMI-1640 with 10% FBS (no cytokines) was added to the cuvette, and the T cells were rested for an additional 60 minutes in the tissue culture incubator. T cells were then maintained at ∼1-1.5e6 cells/mL in T cell media and the media was exchanged every 2-3 days. On day 4-8 post electroporation, engineered T cells were enriched using via NGFR bead selection with the EasySep Release Human PSC-Derived Neural Crest Cell Positive Selection Kit (StemCell Technologies, #100-0047). T cells were used in killing assays a minimum of 10 days after electroporation, and receptor expression and cell purity was assessed via flow cytometry less than 24 hours prior to co-culture with cancer cells.

Jurkat cells (ATCC, #TIB-152) are derived from a T-cell cancer and were engineered to express MANTIS receptors were engineered in a similar way to primary T cells, except that the Jurkat T cells were not bead activated. The electroporation process was kept the same except that 3-5e6 Jurkat cells were resuspended in 100 uL of Lonza SE Buffer (Lonza, #VXC-1024) and electroporated with pulse code CL-120.

### Target Cell Line Generation

NALM6 cell lines virally transduced with GFP-luciferase were generously provided by Martin Pomper^77^, and were cultured in RPMI-1640 (ATCC, #30-2001), 10% Fetal Bovine Serum (FBS, Gibco, # A5670701), and 1% penicillin-streptomycin (P/S, Gibco, # 15140122). To knock-out HLA-A2, NALM6 cells were electroporated with a CRISPR-Cas9 enzyme targeted to the *HLA-A* locus. Briefly, *HLA-A* targeted Cas9 sgRNA (GCTGCGACGTGGGGTCGGAC, IDT) and Cas9 electroporation enhancer (IDT, #1075916 were resuspended at 100 μM concentration in nuclease-free duplex buffer (IDT, #11-01-03-01). CRISPR ribonucleoprotein (RNP) complex was prepared by incubating 50 pmol S.p. Cas9 Nuclease V3 (IDT, # 1081059) with 2-fold excess *HLA-A* sgRNA and 1.5-fold excess Cas9 electroporation enhancer for 15 minutes at room temperature. Then 5e5 NALM6 cells in 20 uL of Opti-MEM were electroporated with CRISPR RNP in a 0.1 cm cuvette (Biorad, #1652089) at 100V for 10 ms using an ECM 2001 Electro Cell Manipulator (BTX). After 7 days of recovery, residual HLA-A2 positive cells were removed from the KO mixture by bead selection with the EasySep Biotin Positive Selection Kit II (StemCell, #17683). For this protocol, a biotinylated anti-HLA-A2 antibody was used (BB7.2, Biolegend, #343322) and the non-bead bound cells were collected. CD19 KO NALM6 were prepared as a part of a previous study^78^ in a similar manner using a *CD19* sgRNA (CGAGGAACCTCTAGTGGTGA).

SKOV3 (ATCC, #HTB-77) cells were cultured in McCoy’s 5a (Gibco, #16600082) supplemented with 10% FBS and 1% PS. SKOV3-Luc cells were used for in vivo studies (Cellomics, #SC-1555). For time-lapse imaging with Incucyte, SKOV3 cells were transduced with Incucyte NucLite Green Lentivirus (nuclear GFP, Sartorius, #4624) in the presence of 5 ug/mL of polybrene and subsequently selected with puromycin. The NucLite GFP SKOV3 was used as the parent cell line for producing EGFR and HER2 KOs. For knocking out EGFR, a cocktail of 3 guide RNAs was used (Hs.Cas9.EGFR.1.AA: TGCTGACTATGTCCCGCCAC, Hs.Cas9.EGFR.1.AB: CCTTGCACGTGGCTTCGTCT, and Hs.Cas9.EGFR.1.AC: GAGTAACAAGCTCACGCAGT). For knock-out HER2, a cocktail of 3 guide RNAs was used: (Hs.Cas9.ERBB2.1.AA: CAACTACCTTTCTACGGACG, Hs.Cas9.ERBB2.1.AB: GCCCTTACACATCGGAGAAC, Hs.Cas9.ERBB2.1.AC: GATAGACACCAACCGCTCTC). For knock-out, 1e6 SKOV3 cells resuspended in 100 μL of Lonza SF Buffer (Lonza, #V4XC-2024) were electroporated with 50 pmol of each Cas9 RNP (150 pmol total RNP) prepared as above with pulse code FE-132 on the Lonza 4D Nucleofector X-Unit (Lonza, AAF-1003X). After electroporation, the cells were rested for 5 minutes at room temperature, and then 500 uL of media was added and the cells were rested for an additional 20 minutes at 37C in an incubator. The cells were then recovered in a T75 flask. KO efficiency was verified by flow cytometry and target null cell lines were isolated via the modified Biotin selection reported above. HER2 KOs were enriched using a biotinylated anti-Her2 antibody (Biotechne, # FAB9589B) and EGFR KOs were enriched using a biotinylated anti-EGFR antibody (Biolegend, #352934). Following enrichment, cells were single cell cloned, and individual clones were used for all *in vitro* experiments. Single cell clones were verified via flow staining and sequencing. A complete EGFR KO line could be generated, but a complete HER2 KO line could not be readily generated possibly due to high dependency of SKOV3 on HER2^58,59^. The SKOV3 HER2 pKO line contained 2/4 functional alleles with in-frame deletions, and it might have expressed a low (but flow undetectable) level of HER2 on the cell surface.

GFP-Luciferized Jurkat Clone E6-1 (ATCC, #TIB-152) cells generated in a previous study^78^ were cultured in RPMI-1640 supplemented with 10% FBS and 1% PS. To overexpress EGFR in Jurkat cells, 1e6 Jurkat cells were transduced with Human EGFR Lentivirus (G&P Biosciences, #LTV0169) in the presence of 8 μg/mL of polybrene overnight. The following day, cells were removed from transduction and allowed to recover for 1 week. EGFR expressing cells were enriched using the EasySep Biotin Positive Selection Kit II (StemCell, #17683) with a biotinylated anti-EGFR antibody (Biolegend, #352934). To create Jurkat cells overexpressing HER2, a truncated variant of HER2 (AA1-684) was knocked-into the TRAC locus using the knock-in approach described above. These cells were also enriched using EasySep Biotin Positive Selection Kit II (StemCell, #17683) except with a biotinylated anti-Her2 antibody (Biotechne, # FAB9589B). Jurkat cells expressing both EGFR and HER2 were prepared similarly, except the truncated HER2 gene was knocked into TRAC locus of the enriched Jurkat EGFR-expressing pool. An EGFR+/HER2+ single cell clone was used *in vivo* experiments.

### Flow Cytometry

Cells were briefly washed and resuspended in PBS by centrifugation. Cell staining was performed at room temperature, except for samples recovered from mice which were stained on ice. Prior to antibody staining, cells were stained with LIVE/DEAD Fixable Near-IR Dead Cell Stain (1 uL/100 uL/1e6 cells; Invitrogen, # L34976) in PBS at room temperature for 10 minutes. Then cells were washed twice in flow staining buffer, which comprised of PBS supplemented with 0.5% bovine serum albumin, 2 mM EDTA, and 0.1% sodium azide. When needed, Fc receptors were first blocked using TruStain FcX PLUS (Biolegend, #156604) and/or Human TruStain FcX (Biolegend, #422302) in 50 μL of flow staining buffer for 10 minutes. Then the desired antibody cocktails were added for 25 minutes for staining in the dark in flow staining buffer. When antibodies with multiple brilliant violet dyes were used, the BD Horizon Brilliant Stain Buffer (BD Biosciences, #566349) was employed. After staining, cells were washed twice with flow staining buffer and samples were run on an Attune NxT flow cytometer (ThermoFisher Scientific). Spectral compensation was conducted using an AbC Total Antibody Compensation Bead Kit (Invitrogen, #A10497) and an ArC Amine Reactive Compensation Bead Kit (Invitrogen, #A10346). Flow data was plotted using FloJo (BD Biosciences, v10.9.0), but spectral compensation was applied to flow data on the Attune NxT Flow Cytometer Software (ThermoFisher Scientific, v6.2.1). Representative gating schemes are shown in Supplementary Figure S1 and S2.

The following antibodies were used for flow cytometry: Brilliant Violet 605™ anti-human CD3 Antibody (BioLegend, 344836), PE/Cyanine7 anti-human CD271 (NGFR) Antibody (BioLegend, 345110), Brilliant Violet 421™ anti-human CD4 Antibody (BioLegend, 300532), PE anti-DYKDDDDK Tag Antibody (BioLegend, 637310), Myc-Tag (9B11) Mouse mAb (Alexa Fluor® 647 Conjugate) (Cell Signaling Technologies, 2233), PE anti-human CD69 Antibody (BioLegend, 310906), Brilliant Violet 421™ anti-human CD69 Antibody (BioLegend, 310930), PE anti-human CD19 Antibody (BioLegend, 302254), APC anti-human HLA-A2 Antibody (BioLegend, 343308), Brilliant Violet 421™ anti-human CD45 Antibody (BioLegend, 304032), Brilliant Violet 605™ anti-mouse CD45 Antibody (BioLegend, 103140), APC anti-human EGFR Antibody (BioLegend, 352906), and the Brilliant Violet 421™ anti-human CD340 (erbB2/HER-2) Antibody (BioLegend, 324420).

### Immunoprecipitation-Flow Cytometry Assay

For the IP-FCM assay, 150,000 Jurkat T cells were co-cultured with 4 μL of anti-c-myc magnetic bead conjugate (Cell Signaling Technologies, #5698) in 200 μL volume in a flat-bottom 96 well plate. Prior to co-culture, beads were washed 3x with cell medium. After 4 hours of culture in an incubator, the magnetic beads were isolated from cells using a 96 well plate magnet. The beads were washed, and residual cells were removed by washing 3x with 200 μL of PBS. The beads were subsequently stained with 100 μL of 1:50 dilution of each antibody for 25 minutes at room temperature. The beads were then washed 3x and then staining was analyzed with flow cytometry.

### T cell Co-Culture Assays

Engineered T cells were incubated with target cells in the amounts and durations listed in the figure legends. Co-cultures with an Incucyte or flow cytometry readout (CD69 flow, SNIPR activation assays, mixed co-culture killing) were setup in 200 μL cytokine free RPMI-1640 medium in 96 well flat bottom plates (Corning, #3595). Flow cytometry was conducted as mentioned above. Interferon gamma from co-cultures was assessed by aspirating 100 μL of the co-culture supernatant and assaying with the Human IFN-gamma ELISA kit (R&D Systems, #DIF50C) according to manufacturer recommended protocols. Live cell imaging of co-cultures was performed with an Incucyte SX5 imaging system (Sartorius). For Incucyte assays, target cells were pre-plated atleast 30 minutes prior to T cell addition. When non-adherent NALM6 target cells were used in incucyte assays, the 96 well plates were coated with 50 μL of 0.1 mg/mL poly-D-lysine (Gibco, #A3890401) solution at room temperature for 60 minutes. The well was then briefly rinsed with 100 μL of sterile water and then air dried for 60 minutes prior to target cell addition. When growth and killing was assessed by a luminescence readout, co-cultures were prepared in 100 μL of cytokine free RPMI-1640 in opaque 96-well flat bottom plates (ThermoFisher Scientific, # 136102). After the culture period, relative target cell viability was assessed using the Steady Glo Luciferase Assay System (Promega, #E2510) using the manufacturer’s recommended protocols. Luminescence was quantified using a Synergy H1 microplate reader (Biotek) with the Gen5 software (Biotek, v2.07).

### Animal Experiments

6–12-week-old female NOD.Cg-*Prkdc^scid^Il2rg^tm1Wjl^*/Szj (NSG) mice were procured either from The Jackson Laboratory (#005557) or from the animal resource facility at The Johns Hopkins Sidney Kimmel Comprehensive Cancer Center. NSG mice were housed in 50% humidity at 70-75°F in a 12-hour light/dark cycle and maintained according to the JHU Animal Care and Use Committee-approved research protocol number MO21M43.

For the mixed NALM6 target cell model, 100k NALM6 Parental (A2+/CD19+), 100k NALM6 A2 KO (CD19+), and 400k NALM9 CD19 KO (A2+) cell lines were injected via tail vein in 200 μL of PBS. Three days after target cell injection, mice were randomized into different groups prior to treatment. 10e6 CAR or MANTIS receptor T cells were injected via tail vein with all cells being normalized 90% NGFR positivity (11.1e6 total T cells). Cells were diluted to similar editing rates by addition of Mock (TCR KO) cells. For the Mock control, 11.1e6 total T cells were injected. The individual performing the injections was blinded to the identity of the treatment groups. On day 10 after T cell injection, 100 μL of blood was collected via submandibular vein bleed into EDTA-treated tubes (Sarstedt, #NC9299309). Red blood cells were then lysed with Ack Lysing buffer (Quality Biological, #118-156-101), and cells were stained using the flow cytometry methods described above. On day 14, mice were sacrificed and the spleens and left femurs were dissected from the mice. To extract the bone marrow, femurs were aspirated with 10 mL of PBS with a 25G syringe into a 70 μL cell strainer (Celltreat, #229483). Spleens or bone marrow aspirate were converted into cell suspensions by mashing with a pestle (Celltreat, #229480) into the cell strainer and rinsing with 10 mL of PBS. Samples were then processed in a similar fashion to peripheral blood samples. To quantify the number of engrafted cells and normalize across different sample sizes, it was assumed that T cell and/or target cell engraftment did not affect the survival of normal murine CD45+ cells in the tissues of interest. Thus, we normalized the count of each cell type based on the murine CD45+ count. To make this adjustment, we multiped each target cell count by the scaling factor defined as:

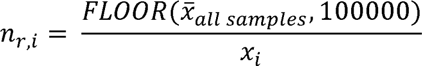

Where *n_r,i_* is the scaling factor for sample *i*, *x_all samples_* is the mean murine CD45+ count across all samples, and *x_i_* is the murine CD45 count in sample *i*. The *FLOOR()* function rounds down the mean murine CD45+ count across all samples to the nearest 1e5. The mean number of murine CD45+ events sampled was ∼5e5 for bone marrow samples and ∼1e5 for splenocytes. Based on this scaling factor, the adjusted target cell count (c_adj,i_) can be calculated from the observed target cell count (*c_obs,i_*) as:

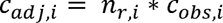

For the orthotopic model of ovarian peritoneal carcinomatosis with HER2 on-target off-tumor toxicity, 4e6 SKOV3-Luc cells were intraperitoneally injected in 200 μL of PBS. Ten days after inoculation, tumor engraftment was verified using IVIS bioluminescence imaging (BLI) and mice were randomized into groups. The experimenter was blinded to the identity of all treatment groups. For BLI imaging, mice were i.p. injected with 150 μL of IVISbrite D-Luciferin Ultra Bioluminescent Substrate in RediJect Solution (Revvity, #770505) and then anesthetized with inhaled isoflurane. Mice were imaged 9-10 minutes post injection using an IVIS spectrum imager (Revvity). Tumor burden was quantified by integrating total luminescence signal across captured images for each mouse with the Living Image software (Revvity, v.4.5.5). On day 11, 14e6 CAR T cells at 90% NGFR positivity with Mock TCR KO T cells (∼15.5e6 total T cells) were i.p. injected in 200 μL of PBS. CAR T cells with higher purity were normalized to this NGFR percentage by dilution with Mock (TCR KO) T cells. Following treatment, mice were closely monitored for signs of toxicity and were euthanized if they showed greater than 20% weight loss not recovering within 24 hours or other signs of distress that included a combination of hunched posture, labored breathing, ruffled coat, lethargic activity, hind-leg paralysis, or significant edema with veterinary recommendation. CAR T cell engraftment and BLI were monitored routinely as described above. To assess toxicity in mice, necropsies were performed by extracting organs, fixing and paraffin embedding tissues, and preparing H&E-stained slides which were reviewed by a veterinary pathologist.

For the Jurkat EGFR+/HER2+ cell-line derived xenograft model, 1e6 cancer cells were intravenously injected into mice in 200 μL of PBS. IVIS imaging was performed as described above 1 hour after injection, and mice were randomized into groups based on this measurement. The experimenter was blinded to all groups. Two days later, on Day 12 post T cell engineering, mice were treated with 10e6 CAR T cells normalized to 80% NGFR positivity via addition of Mock (TCR KO) T cells (∼12.5e6 total T cells) via i.v. injection in 200 μL of PBS. Tumor burden was quantified over time based on the average radiance signal across the whole mouse body. Following treatment, mice were closely monitored for signs of toxicity and were euthanized if they showed greater than 20% weight loss not recovering within 24 hours or other signs of distress including a combination of hunched posture, labored breathing, ruffled coat, lethargic activity, hind-leg paralysis, or significant edema with veterinary recommendation. Mice showing signs of toxicity and/or weight loss greater than 15% were subcutaneously injected daily with 1-2 mL of normal saline to prevent dehydration until weight recovered, symptoms resolved, or the mice met the criteria for euthanasia described above.

### Statistical Analysis & Plotting

Statistical analysis was performed with Prism v10.2.1 (Graphpad) or in python with the lifelines package (v.0.30.3). The statistical test that was used for each analysis is indicated in the figure legends. Some plots were generated using Prism v5.00 (Graphpad). All graphics were generated with protein structures rendered via ChimeraX (UCSF, v1.8) and then converted into bitmap vector images with Inkscape (v1.4). Graphical representations depicting cells and mice were modified from images deposited on NIH BioArt.

## Acknowledgements

We would like to thank Kathleen Helwig, Leslie Wang, Ali Dbouk, Alisha Mills, and Ilene Vogelstein for administrative support on this project. In the Kinzler/Vogelstein lab, we would like to acknowledge current and former members: Michael Hwang, Nicholas Wyhs, Xuyang Li, Tolulope Awosika, Stephanie Glavaris, Jiaxin Ge, Joshua Urban, Jacqueline Douglass, Yuanxuan Xia, Sze Kiat Tan, Connor Liu, Bracha Erlanger Avigdor, Zaid Bayyat, Raghad Al Ishaq, Akash Jain, Riana Sanjeevan, Noah Brookes, Raygan Murray, Adaira Dumm, Ashley Djuhadi, Nihao Sun, Petvy Li, and Samuel Curtis for helpful scientific discussion. We would also like to acknowledge Maximilian Konig and members of his lab including Jin Liu, Colin Gliech, Brock Moritz, and Kyle Kaeo for helpful scientific discussion. We would like to acknowledge Tian-Li Wang and Ellen Tully for generously sharing the SKOV3 sublines they generated (not used in this study) and their experience with the SKOV3 cells. Finally, from the BKI Flow Core, we would like to acknowledge Ada Tam for her support on flow sorting.

## Funding

The Virginia and D.K. Ludwig Fund for Cancer Research

Lustgarten Foundation for Pancreatic Cancer Research

The Commonwealth Fund

The Bloomberg Kimmel Institute for Cancer Immunotherapy

Bloomberg Philanthropies

NIH Cancer Center Support Grant P30 CA006973

TDN was supported by NCI grant T32 CA153952

BJM, SRD & AHP were supported by NIH Grant T32 GM136577

NM was supposed by grant NIGMS T32 GM148383

SP was supported by NCI Grant K08CA270403, the Blood Cancer United Translation Research Program award, the American Society of Hematology Scholar award, and the Swim Across America Translational Cancer Research Award

CB was supported by NCI Grant R37 CA230400

## Author contributions

Conceptualization: TDN, BJM, AHP, BV, SZ, KWK

Methodology: TDN, BJM, AHP, SRD, MDM, SS, SP, KG, EW

Investigation (*in vitro*): TDN, TSA, NM

Investigation (animal experiments): TDN, EW, RMC, KG

Analysis and interpretation of data: TDN, BJM, AHP, TSA, NM, RMC, EW, BSL, SRD, NP, CB, MDM, DW, DMP, SP, SS, KG, SZ, BV, KWK

Writing: original draft: TDN

Writing: review & editing: TDN, SZ, BV, KWK, SP, AHP, SS, DMP

Supervision: KWK, SZ, BV

## Competing interests

BV & KWK are founders of Exact Sciences. KWK and NP are advisors to, and hold equity, in Exact Sciences. BV, KWK, and NP are founders of, and own equity in, Haystack Oncology, a Quest Diagnostics company, as well as Clasp Therapeutics and Winnow Therapeutics. NP is a consultant to Vidium. BV is a consultant to and holds equity in Catalio Capital Management. S.Z. owns equity in Exact Sciences, is a founder of, holds equity in, and serves as consultants to Clasp Therapeutics and Winnow Therapeutics, and is also a consultant to and holds equity in NeoPhore. SZ has a research agreement with BioMed Valley Discoveries, Inc. CB is a consultant to Haystack Oncology, Privo Technologies and Longeviti Neuro Solutions. CB was a co-founder of OrisDx. CB is a co-founder of Belay Diagnostics. DMP reports grant and patent royalties through institution from BMS, a grant from Compugen, stock from Trieza Therapeutics and Dracen Pharmaceuticals, and founder equity from Potenza; being a consultant for Aduro Biotech, Amgen, Astra Zeneca (Medimmune/Amplimmune), Bayer, DNAtrix, Dynavax Technologies Corporation, Ervaxx, FLX Bio, Rock Springs Capital, Janssen, Merck, Tizona, and Immunomic-Therapeutics; being on the scientific advisory board of Five Prime Therapeutics, Camden Nexus II, WindMil; being on the board of directors for Dracen Pharmaceuticals. S.P. is a consultant to Merck, Nexoplex Bio, owns equity in Gilead and received payments from IQVIA and Curio Science. S.P. is a founder and hold equity in Winnow Therapeutics. S.S. is a consultant and holds equity in Cage Pharma Inc. The Johns Hopkins University has filed patent applications related to technologies described in this paper on which TDN, BM, AHP, SZ, BV, and KWK are listed as inventors. The companies named above, as well as other companies, have licensed previously described technologies related to the work described in this paper from Johns Hopkins University. BV, KWK, SZ, NP are inventors of some of these technologies. Licenses to these technologies are or will be associated with equity or royalty payments to the inventors as well as to Johns Hopkins University. The terms of all these arrangements are being managed by Johns Hopkins University following its conflict of interest policies.

## Data and Materials availability

Plasmids are available by request under a material transfer agreement with Johns Hopkins University.

## Supplementary Materials

### Extended Data Figures

**Extended Data Fig. 1.**
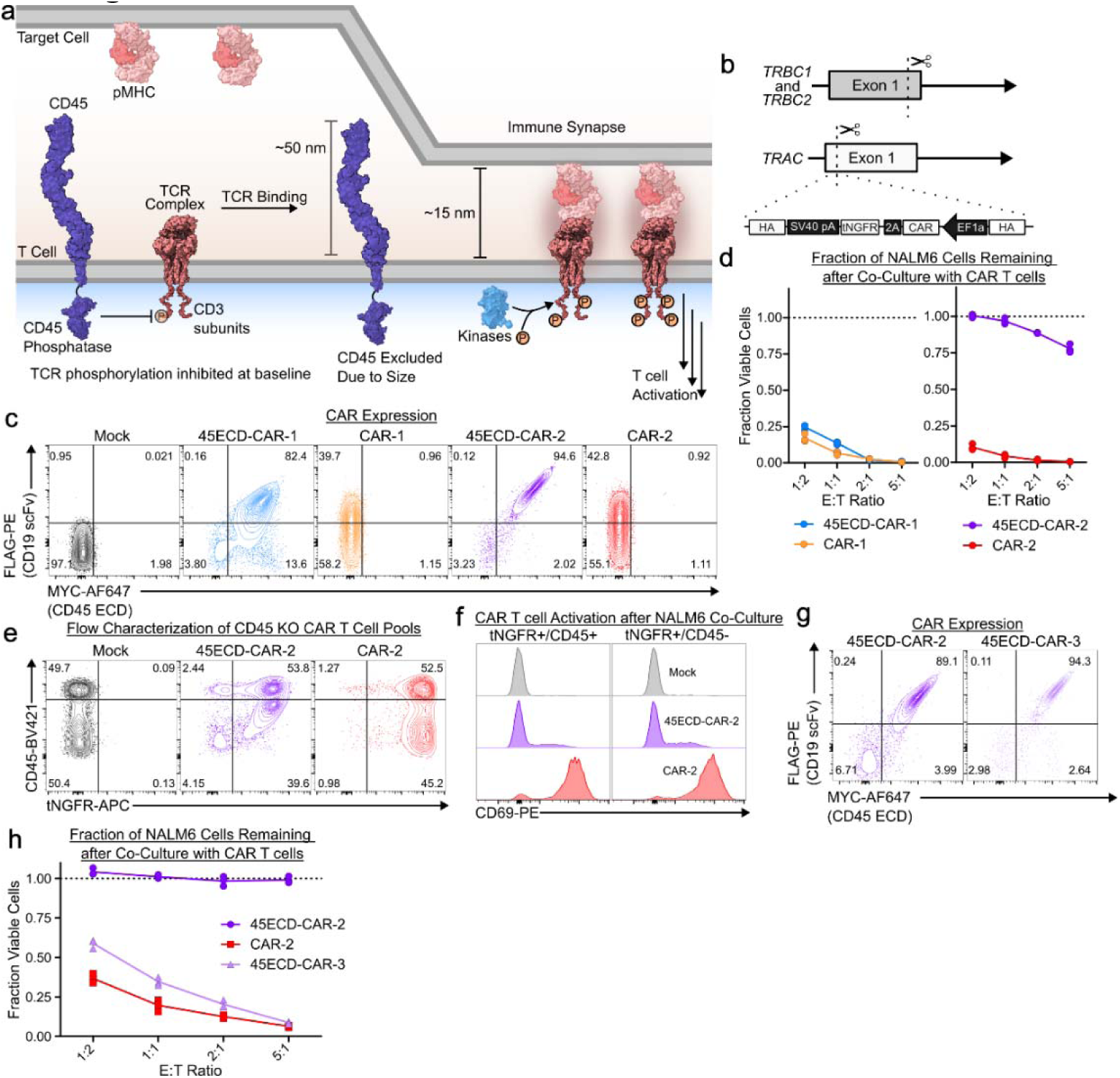
**(a)** A spatial model of T cell signaling. CD45 phosphatase activity is regulated by its large bulky extracellular domain, which cannot physically fit in the immunological synapse. Exclusion from the immune synapse enables efficient TCR phosphorylation. TCR = T cell receptor. pMHC = peptide-Major Histocompatibility Complex. **(b)** CAR transgenes were knocked into the *TRAC* locus using CRISPR-mediated homology directed repair in primary T cells. *TRBC* was simultaneously knocked out to ensure all cells were TCR negative. The CAR transgenes encoded a truncated NGFR (tNGFR) tag for selection of knocked-in T cells via bead selection. This approach was used for all constructs described in this study. **(c)** CARs and 45ECD-CAR chimeras were enriched with NGFR selection and assessed for receptor expression via flow cytometry. The MYC tag is incorporated on the CD45 ECD component and the FLAG tag is incorporated on the CD19 scFv. **(d)** 5,000-50,000 45ECD-CARs and controls were incubated with 10,000 NALM6 GFP-luciferase cells at varying E:T ratios for 48 hours. Cell viability was measured using a SteadyGlo luminescence assay. The fraction of viable cells compared to the Mock condition is depicted. **(e)** CAR T cells were engineered with an additional CRISPR-Cas9 enzyme targeted to endogenous CD45. Flow characterization of the NGFR, but not CD45 selected CAR T cell product. **(f)** 50,000 polyclonal CD45 KO CAR T cells were co-cultured with NALM6 cancer cell lines in a 1:2 E:T ratio for 24 hours and CD69 expression was assessed in the engineered CD45+ and CD45 null lines based on the gates indicated in d. **(g)** Flow cytometry of 45ECD-CAR-2 and 3 constructs as in **b**. **(h)** Cytotoxicity of 45ECD-CAR-2, 45ECD-CAR-3, and CAR-2 at varying E:T ratios in the same way as in **c**. Data in **d, f,** and **h** are representative of n = 3 technical replicates.

**Extended Data Fig. 2.**
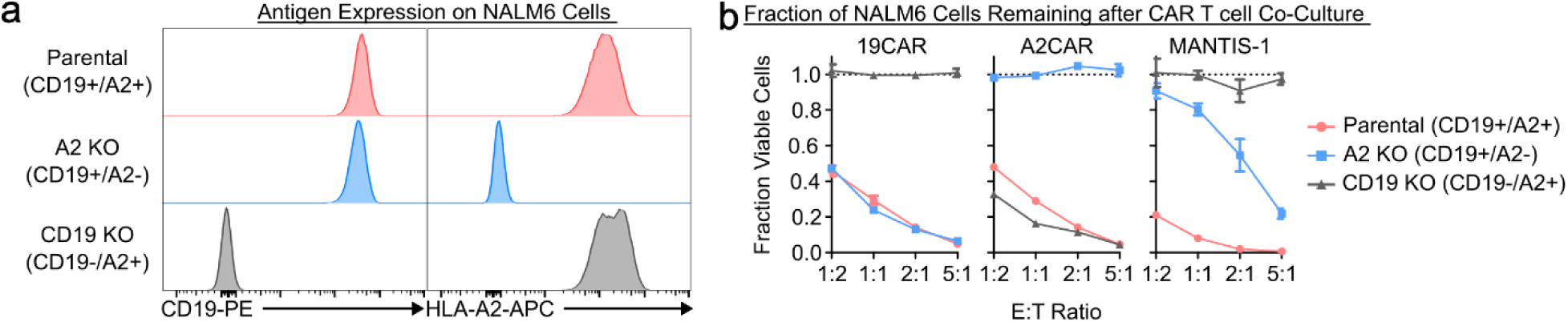
**(a)** Flow staining of NALM6 isogenic lines generated via CRISPR-Cas9 mediated knock-out. NALM6 A2 KO (blue) represents a polyclonal pool prepared by KO and bead selection of HLA-A2 negative cells. CD19 KO pool (grey) is a polyclonal pool generated by KO of CD19 and mixing single cell clones. **(b)** 5,000-50,000 CAR controls and MANTIS cells were co-cultured with 10,000 NALM6 isogenic cell lines at varying E:T ratios. After 48 hours in culture, the fraction of viable cells was determined using a SteadyGlo luminescence assay. The fraction of viable cells compared to the Mock condition is depicted. Data in **b** represents n = 3 technical replicates.

**Extended Data Fig. 3.**
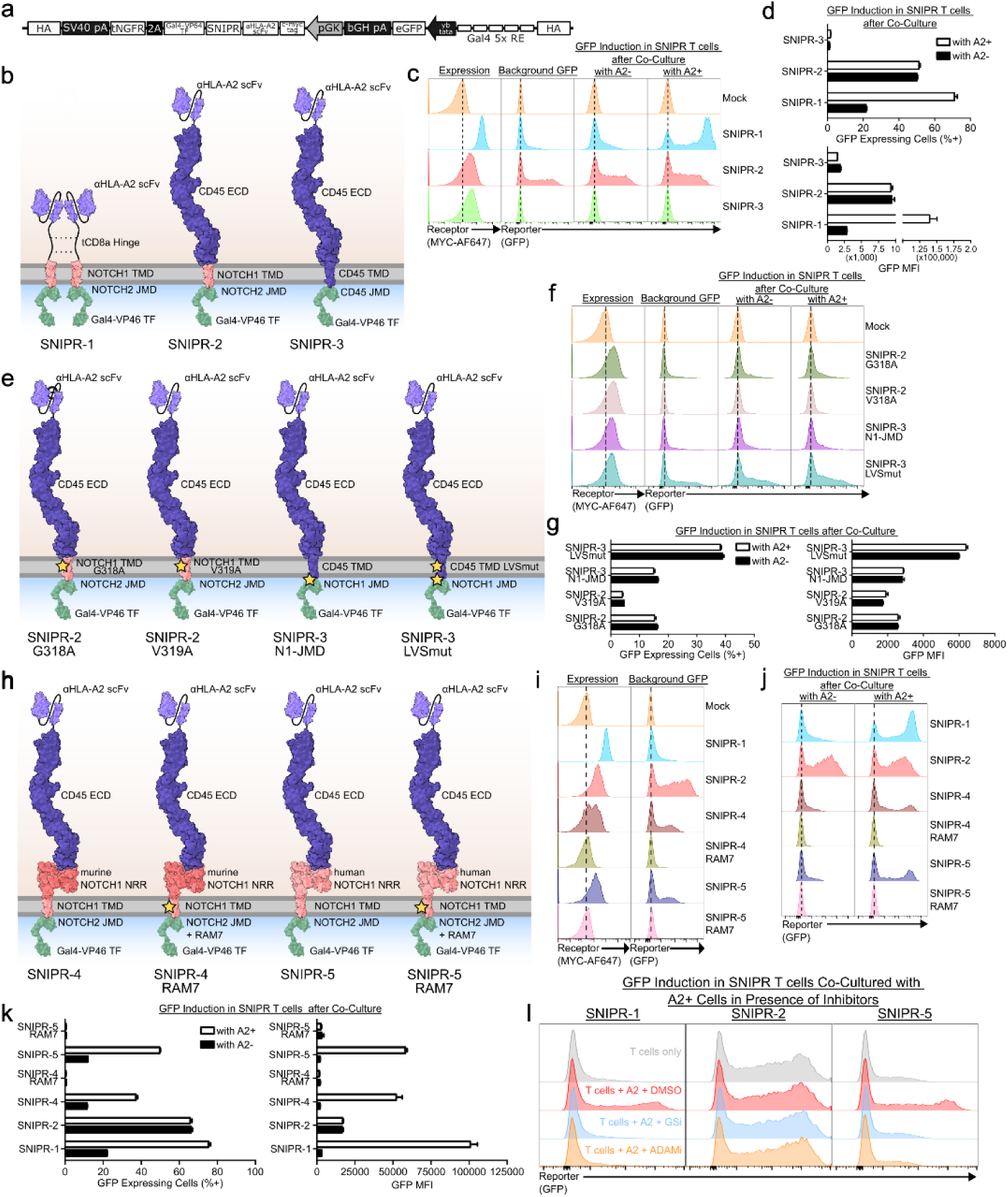
**(a)** Design and construction of HLA-A2 targeted SNIPR transgenes which were knocked-in to the *TRAC* locus of primary T cells. SNIPR expression was mediated by a constitutive pGK promoter. Intracellular release of a Gal4_VP64 transcription factor mediated inducible expression of GFP. **(b)** Design of SNIPR receptors. SNIPR-1 is an optimized SNIPR design previously reported^45^. In SNIPR-2, the tCD8a Hinge of SNIPR-1 was replaced with CD45 ECD derived from component of MANTIS-1. In SNIPR-3, the CD45 TMD/JMD replaced the NOTCH-1 TMD/NOTCH-2 JMD in SNIPR-2. **(c,d)** 50,000 SNIPR T cells were incubated with or without 50,000 NALM6 Parental (A2+) or A2 KO (A2-) target cells for 48 hours. The percent of GFP+ cells and median fluorescence intensity were measured with flow cytometry of SNIPR T cells. **(e)** Design of SNIPR-2 and SNIPR-3 mutants that were expected to reduce or increase reporter activity respectively. G318A and V319A reduce background activity of NOTCH transmembrane domains^45^. Changing the acidic JMD of CD45 to that of NOTCH was expected to enhance proteolysis, as is the addition of an LVS gamma secretase site^45^. **(f,g)** Same as **b,c,** but with the SNIPRs in **e**. **(h)** Design of CD45 ECD SNIPRs with murine and human NRRs and a RAM7 sequence reported to reduce ligand independent activity. **(i,j,k)** Same as in **b,c**, but with the SNIPRs in **h**. Including NRRs creates ligand dependent activity of CD45 ECD SNIPRs. **(l)** SNIPRs were co-cultured in the presence of 10 μM DAPT (gamma secretase inhibitor, GSi) or 25 μM GI254023X (ADAM inhibitor, ADAMi) with A2+ NALM6 cells. After 48 hours, GFP reporter activity was assessed. All data representative of n = 3 technical replicates. The mean and standard deviation is plotted of all replicates in **d**, **g**, and **k.**

**Extended Data Fig. 4.**
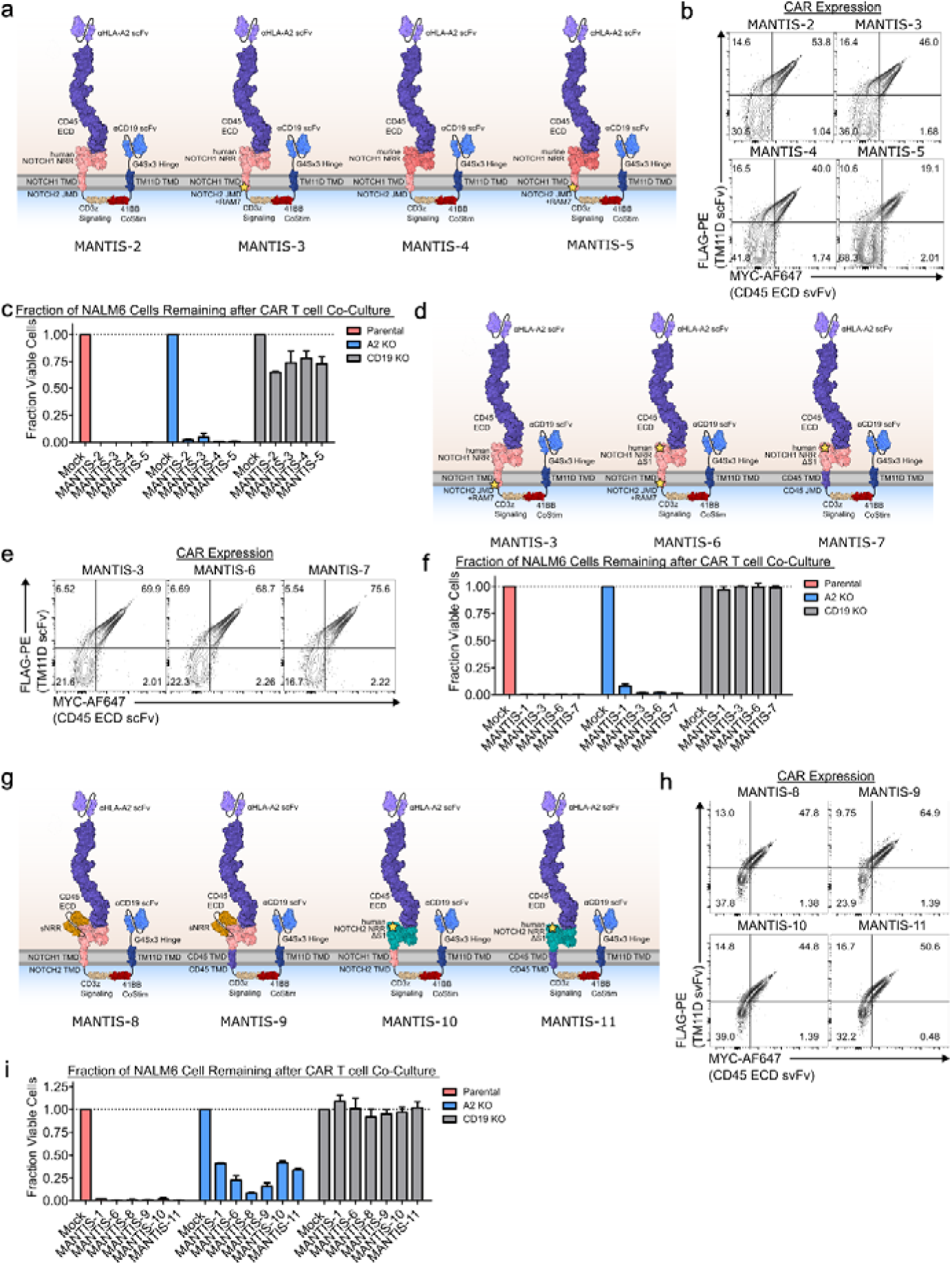
**(a)** MANTIS designs incorporating negative regulatory regions (NRRs). Some variants included a hydrophobic RAM7 peptide^49^. **(b)** Flow staining of MANTIS designs after NGFR selection. The MYC tag is N-terminal to the HLA-A2 scFv, and the FLAG tag is C-terminal on the CD19 scFv. **(c)** 25,000 MANTIS cells in **c** were co-cultured with 25,000 NALM6 isogenic lines: NALM6 Parental (A2+/CD19+), NALM6 A2 KO (CD19+), and NALM6 CD19 KO (A2+). After 72 hours, cell killing was measured via flow cytometry and normalized to counts in the Mock condition. **(d)** MANTIS designs incorporating deletion of the S1 domain of human NRR and incorporating a CD45 TMD/JMD. **(e)** Expression of MANTIS-3, 6, 7 via flow cytometry. **(f)** 50,000 MANTIS cells were co-cultured with 50,000 NALM6 isogenic lines for 48 hours. Cell killing was measured by a SteadyGlo luminescence assay. The fraction of viable cells compared to the Mock condition is depicted. These modifications also did not improve the specificity of MANTIS. **(g)** New MANTIS designs incorporating a sNRR^52^or a human NOTCH2 NRR with CD45 TMD/JMD. **(h)** Expression of MANTIS designs in **g**. **(i)** 50,000 MANTIS cells were co-cultured with 50,000 NALM6 isogenic lines. 24 hours later, remaining target cells were assessed via flow cytometry and counts of each line were normalized to the Mock condition. Data in **c, f,** and **i** are representative of n = 3 technical replicates. The mean and standard deviation is plotted of all replicates.

**Extended Data Fig. 5.**
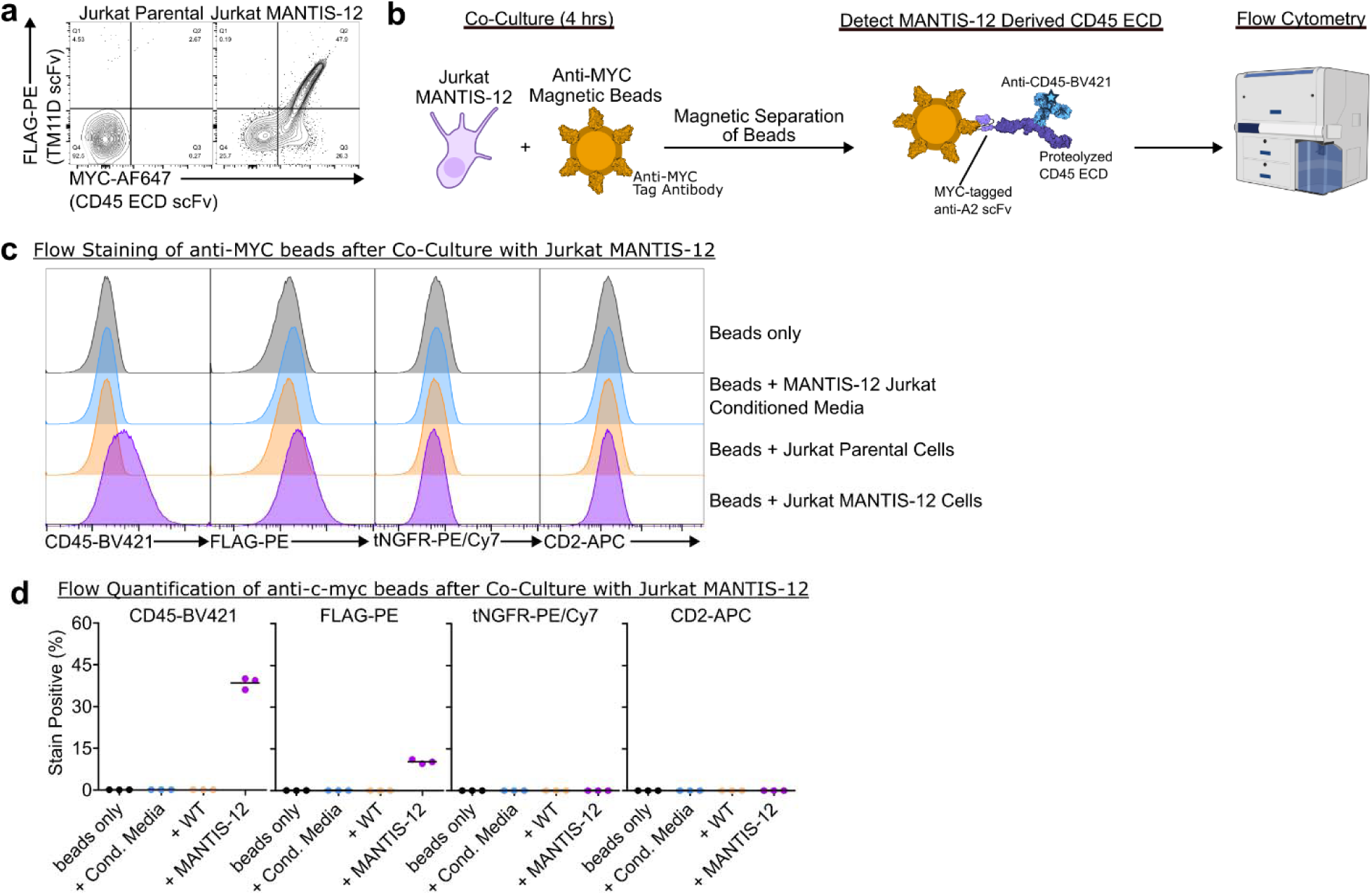
MANTIS-12 was expressed in Jurkat T cells and **(a)** receptor expression was assessed via flow cytometry. The MYC tag is N-terminal to the anti-A2 scFv; the FLAG tag is on C-terminal to the anti-CD19 scFv depicted in Fig. 3a. **(b)** A 1:40 volumetric dilution of Anti-MYC beads were co-cultured with 150,000 Jurkat cells with or without MANTIS-12 receptors for 4 hours to stimulate MYC-tagged MANTIS-12 on Jurkat cells. Beads were also co-cultured with conditioned culture media (Cond. Media) from Jurkat MANTIS-12 to verify absence of non-specific capture of CD45 fragments in culture medium. The beads were then isolated via magnetic separation and stained with various antibodies to detect proteolyzed MANTIS-12 components transferred onto beads. The stained beads were analyzed by flow cytometry. **(c)** Histograms of anti-MYC beads stained with various antibodies after co-culture with Jurkat cells in **a**. **(d)** Quantified percent staining from the histograms depicted in **c**. FLAG tag staining is indicative of capture of full-length MANTIS-12 fragments. Data in **c** and **d** are representative of n = 3 technical replicates. Bar represents the mean of all replicates.

**Extended Data Fig. 6.**
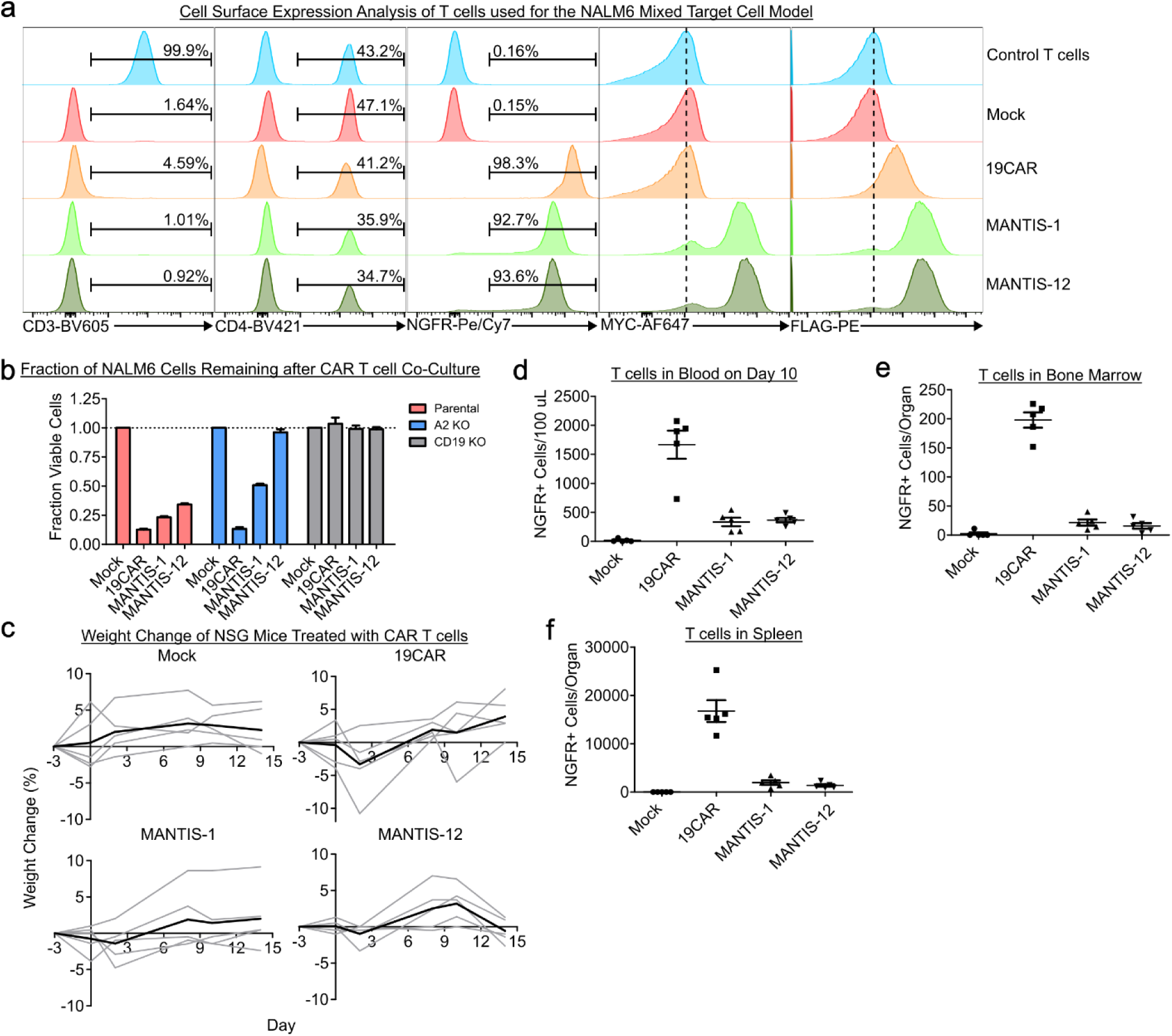
Supporting information for the experiment depicted in Fig. 4d. **(a)** Flow characterization of the T cells which were injected into mice. Control T cells were used as flow staining control only and were not injected into mice. **(b)** For quality control purposes, T cells which were to be injected into mice were co-cultured in a 1:1 E:T ratio with 50,000 NALM6 isogenic lines overnight. Cell viability was assessed via SteadyGlo luminescence. Data represents mean and standard deviation of n = 3 technical replicates. **(c)** Weight change of each individual mice during treatment. Grey lines depict individual mice, and black lines are the average of all mice in each group. **(d**) On day 10, peripheral blood from NSG mice was obtained and cells were stained for NGFR to identify engrafted T cells. Count of T cells identified in **(e)** bone marrow and **(f)** spleens in mice after 14 days of treatment. Each data point in **d, e,** and **f** is representative of one mouse. Bars represent mean +/− standard deviation. There were n = 5 mice per group in this experiment.

**Extended Data Fig. 7.**
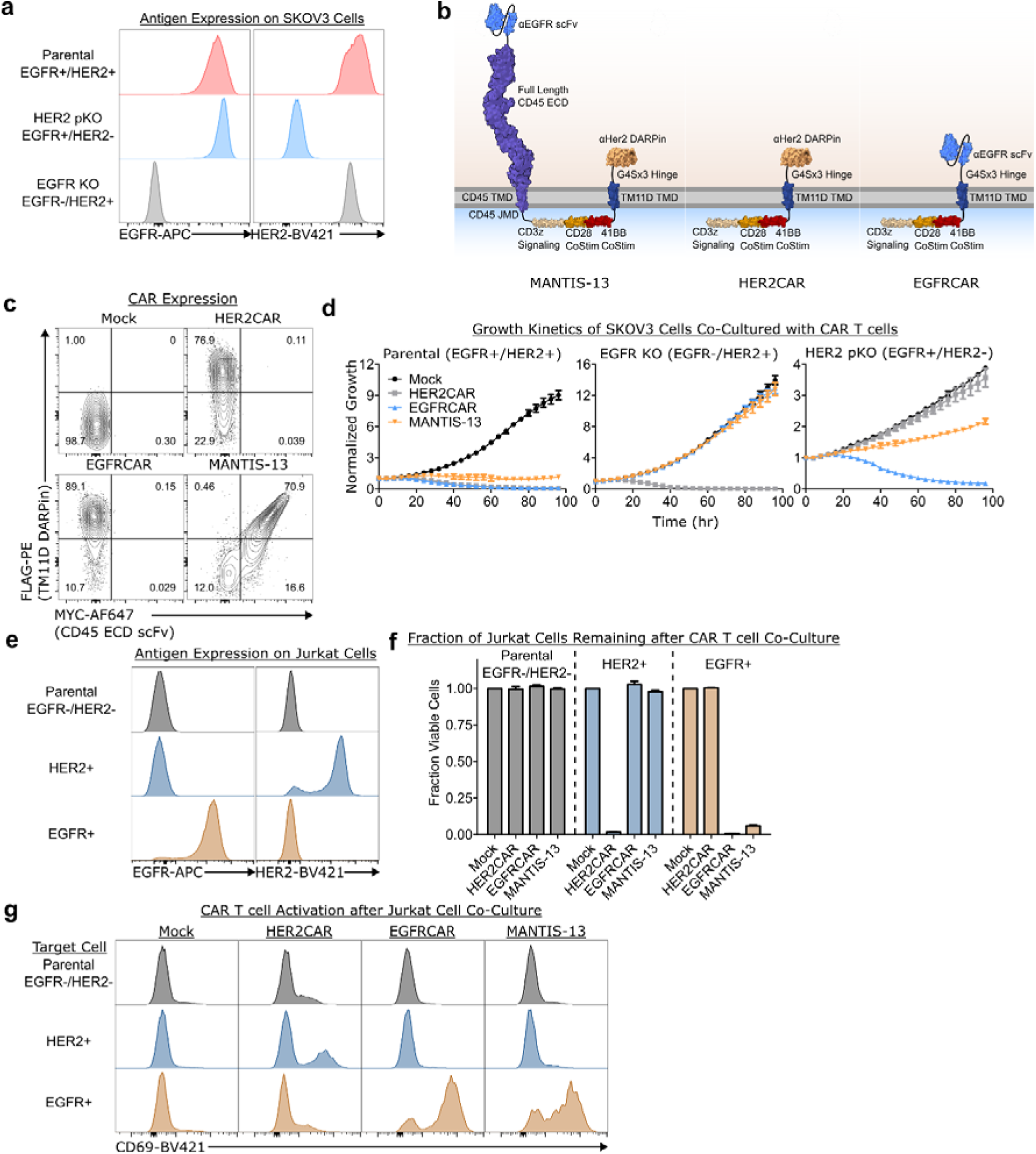
**(a)** Flow staining of SKOV3 isogenic lines generated via CRISPR-Cas9 KO. Note that HER2 was only partially knocked out as noted in the text. **(b)** MANTIS-13 and control CAR construct designs incorporating an additional CD28 co-stimulation domain. **(c)** Expression of MANTIS-13 and controls CARs by flow staining for MYC and FLAG tags placed at the N- and C-terminus of all constructs. CAR controls only encode a FLAG tag. **(d)** 2,000 SKOV3 GFP+ cells were co-cultured with the T cells described in **b** at a 5:1 E:T ratio. Target cell growth was measured with time-lapse imaging. Normalized Growth represents total GFP integrated object intensity normalized by the integrated intensity at the zero hr timepoint. **(e)** Expression of EGFR and HER2 on GFP-luciferase+ Jurkat cells engineered to artificially express these antigens. **(f)** 25,000 Jurkat target cells in **e** were co-cultured with 25,000 MANTIS-13 T cells or control CAR T cells for 48 hours and target cell viability was measured with SteadyGlo luminescence. The fraction of viable cells compared to the Mock condition is depicted. **(g)** CD69 expression on MANTIS-13 and control CAR T cells after co-culture with 25,000 Jurkat target cells in a 1:1 ratio for 24 hours. Data in **d**, **f**, and **g** are representative of n = 3 technical replicates. The mean and standard deviation is plotted in **d** and **f**.

**Extended Data Fig. 8.**
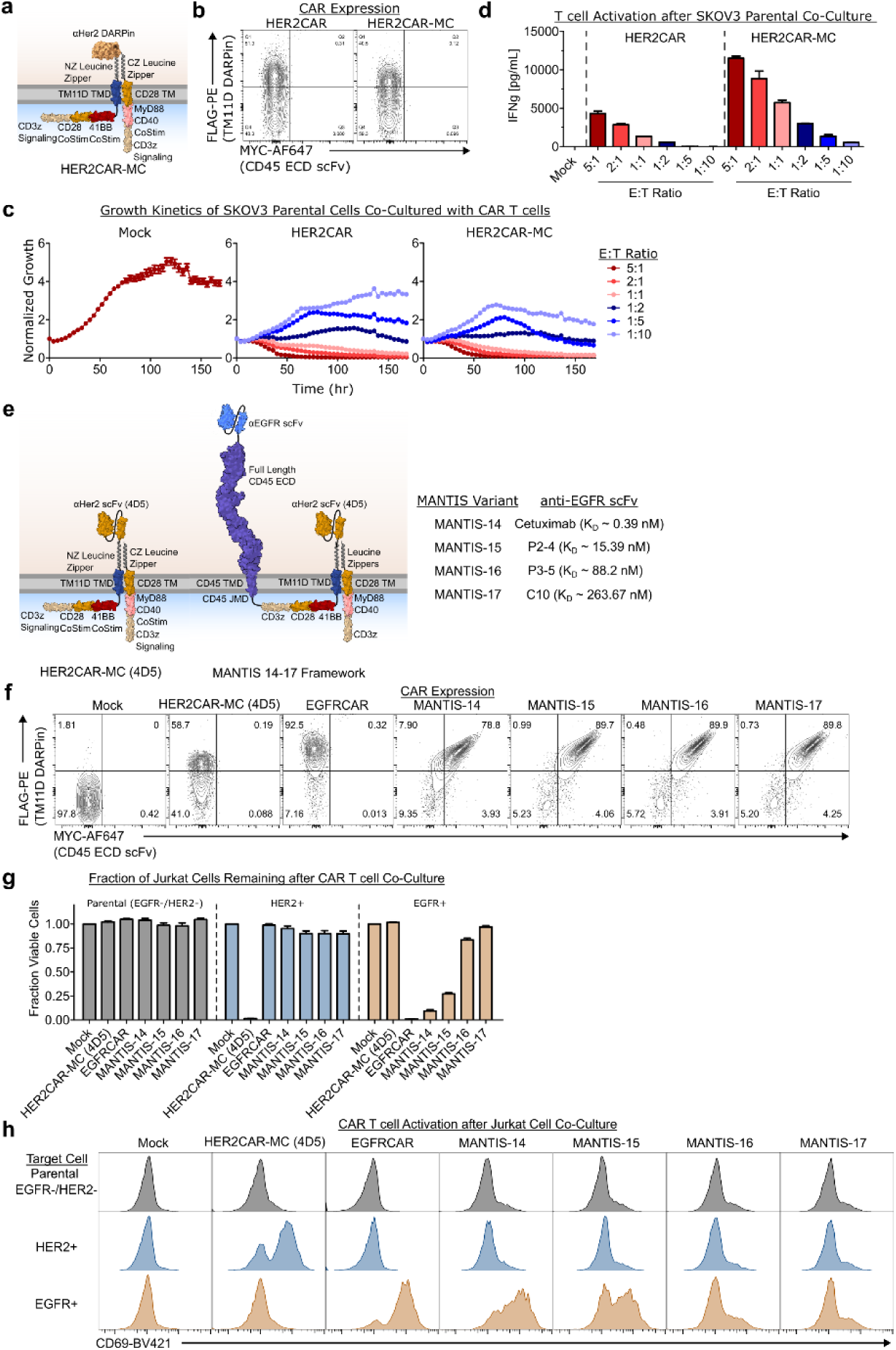
**(a)** Design of HER2CAR-MC, in which an additional MyD88CD40-CD3z domain was appended to the DARPin CAR via anti-parallel NZ and CZ leucine zippers. **(b)** Expression of HER2CAR and HER2CAR-MC was assessed via flow cytometry. The FLAG tag is expressed on the extracellular C-terminus of the DARPin, but no MYC tag is present in these constructs. **(c)** 5,000 SKOV3 WT cells were co-cultured with HER2CAR or HER2CAR-MC at varying E:T ratios. The total number of T cells in the well was kept constant via the addition of non-functional Mock T cells. Normalized Growth represents total GFP integrated object intensity normalized by the integrated intensity at the zero hr timepoint. **(d)** IFNg secretion by CAR T cells from the co-cultures described in **c** at the 72-hour timepoint. **(e)** MANTIS constructs including the MC module. In these constructs, the 4D5 scFv was used instead of the H10-2-G3 DARPin. Different anti-EGFR binders with varying affinities were also used to test the effect on MANTIS specificity. **(f)** Same as **b** but for the constructs depicted in **e**. **(g)** 25,000 Jurkat target cells were co-cultured with 25,000 MANTIS T cells or control CAR T cells for 48 hours. The fraction of viable cells was measured by a SteadyGlo luminescence assay. The fraction of viable cells compared to the Mock condition is depicted. **(h)** CD69 expression on MANTIS-14-17 and control CAR T cells after co-culture with 25,000 Jurkat target cells in a 1:1 ratio for 24 hours. Data in **c-d** and **g-h** are representative of n = 3 technical replicates. The mean and standard deviation is plotted in **d**, **c**, and **g**.

**Extended Data Fig. 9.**
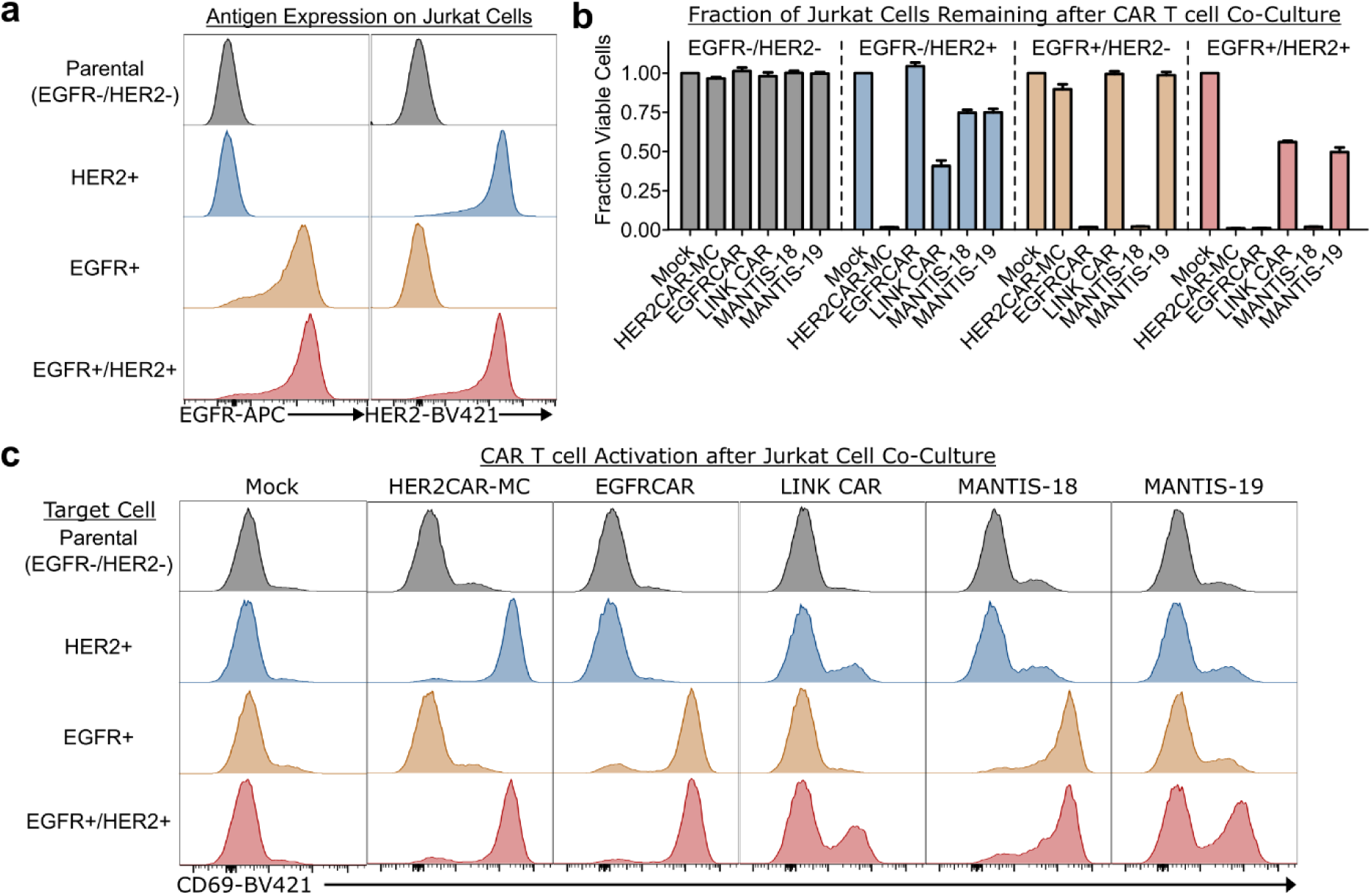
**(a)** Expression of EGFR and HER2 on Jurkat cell lines engineered to express each antigen individually or both antigens simultaneously. **(b)** 25,000 MANTIS, LINK CAR, or control CAR T cells were co-cultured with 25,000 Jurkat cells for 48 hours. The fraction of remaining Jurkat was determined by a SteadyGlo luminescence assay. The fraction of viable cells compared to the Mock condition is depicted. **(c)** CD69 expression on MANTIS 18-19, LINK CAR, or control CAR T cells after co-culture with 25,000 Jurkat target cells in a 1:1 ratio for 24 hours. Data in **b** and **c** are representative of n = 3 technical replicates. The mean and standard deviation is plotted in **b**.

**Extended Data Fig. 10.**
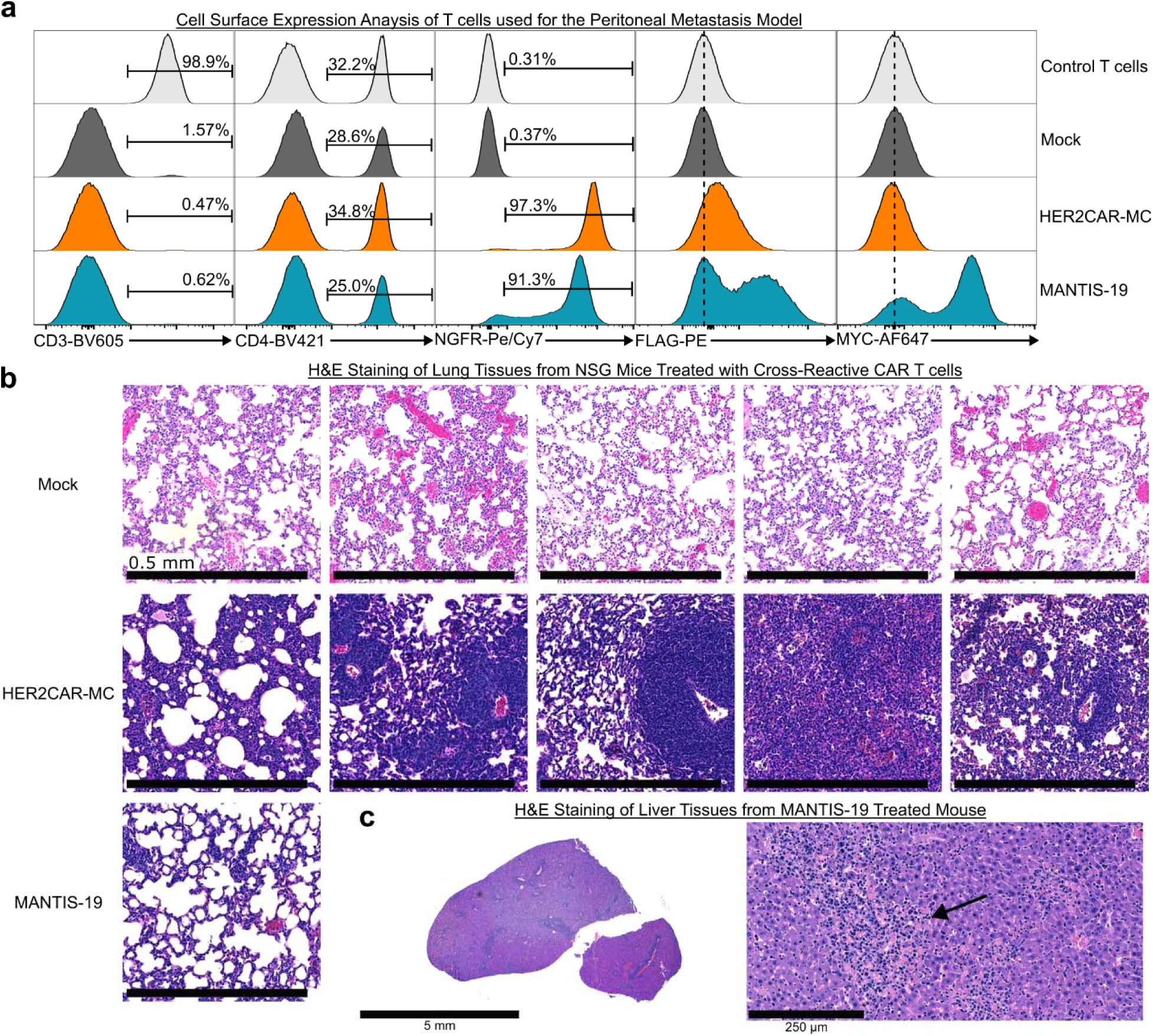
**(a)** Cell surface staining of CAR T cells used in the model of SKOV3 peritoneal carcinomatosis depicted in Fig. 5. Control T cells were not infused into the mice and are depicted as flow staining control. **(b)** H&E staining of representative lung tissue sections from dead animals depicted in Fig. 5. **(c)** H&E staining of liver tissue from the one MANTIS-19 mouse that died early in the experiment. Left, low magnification image. Right, high magnification image with a representative necrotic site, showing immune infiltration (black arrow).

**Extended Data Fig. 11.**
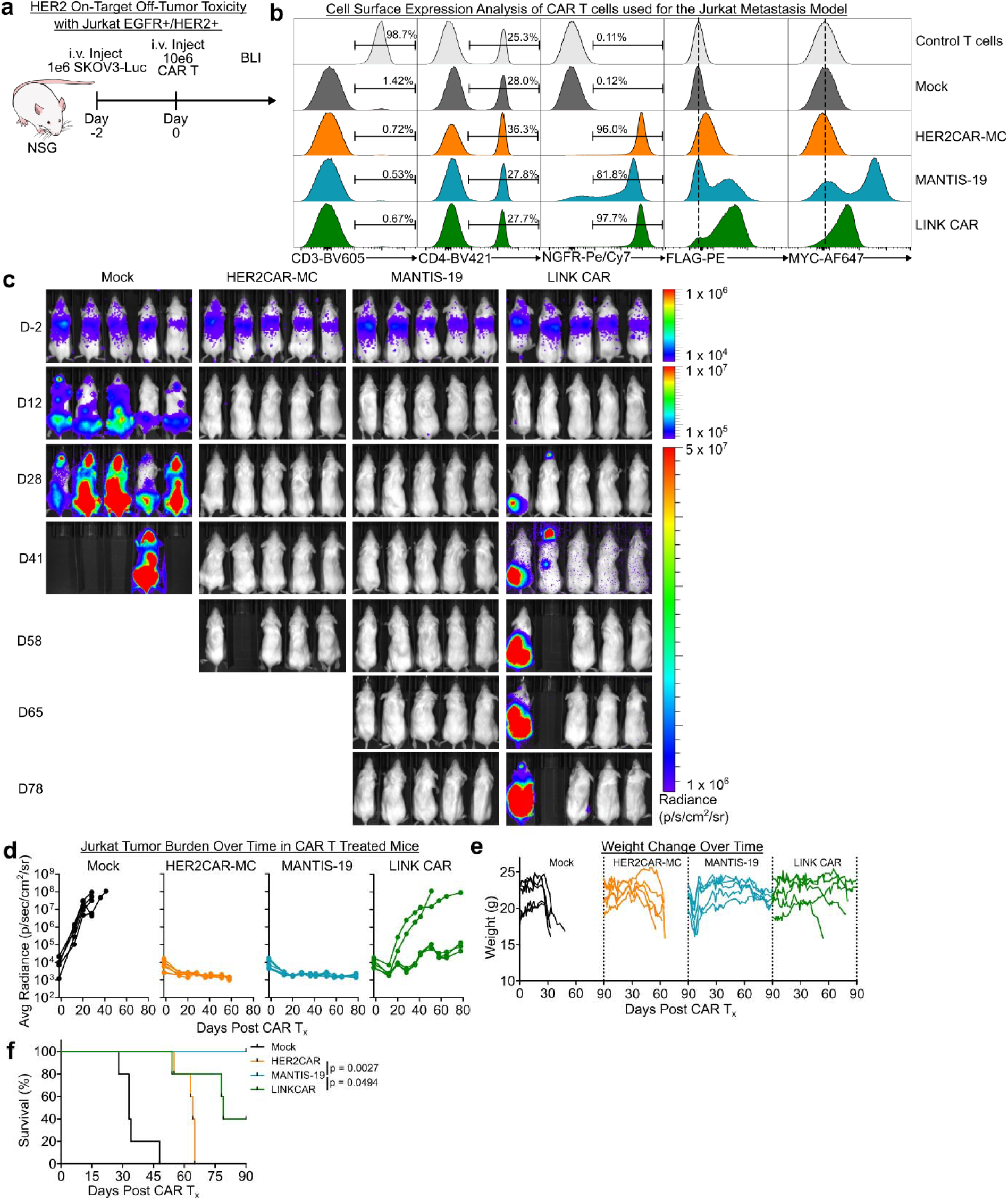
**(a)** Timeline for the NSG mouse experiment to study HER2 on-target off-tumor toxicity with Jurkat EGFR+/HER2 xenografts implanted via tail vein injection. Mice were treated i.v. with 10 million Mock, HER2CAR-ME, MANTIS-19, or LINK CAR T cells. **(b)** Cell surface staining of various T cells used in the mouse model. Control T cells were not infused into mice and were used only as a flow staining control. **(c)** Tumor burden assessed by bioluminescence imaging. **(d)** Quantification of the tumor burden illustrated in **c**. **(e)** Weight change and **(f)** survival of mice after various treatments. p values calculated using a log-rank (Mantil-Cox) test. Data in **c-f** representative of n = 5 mice per group. BLI = bioluminescence imaging. T_x_ = treatment.

### Supplementary Figures

**Supplementary Figure S1.**
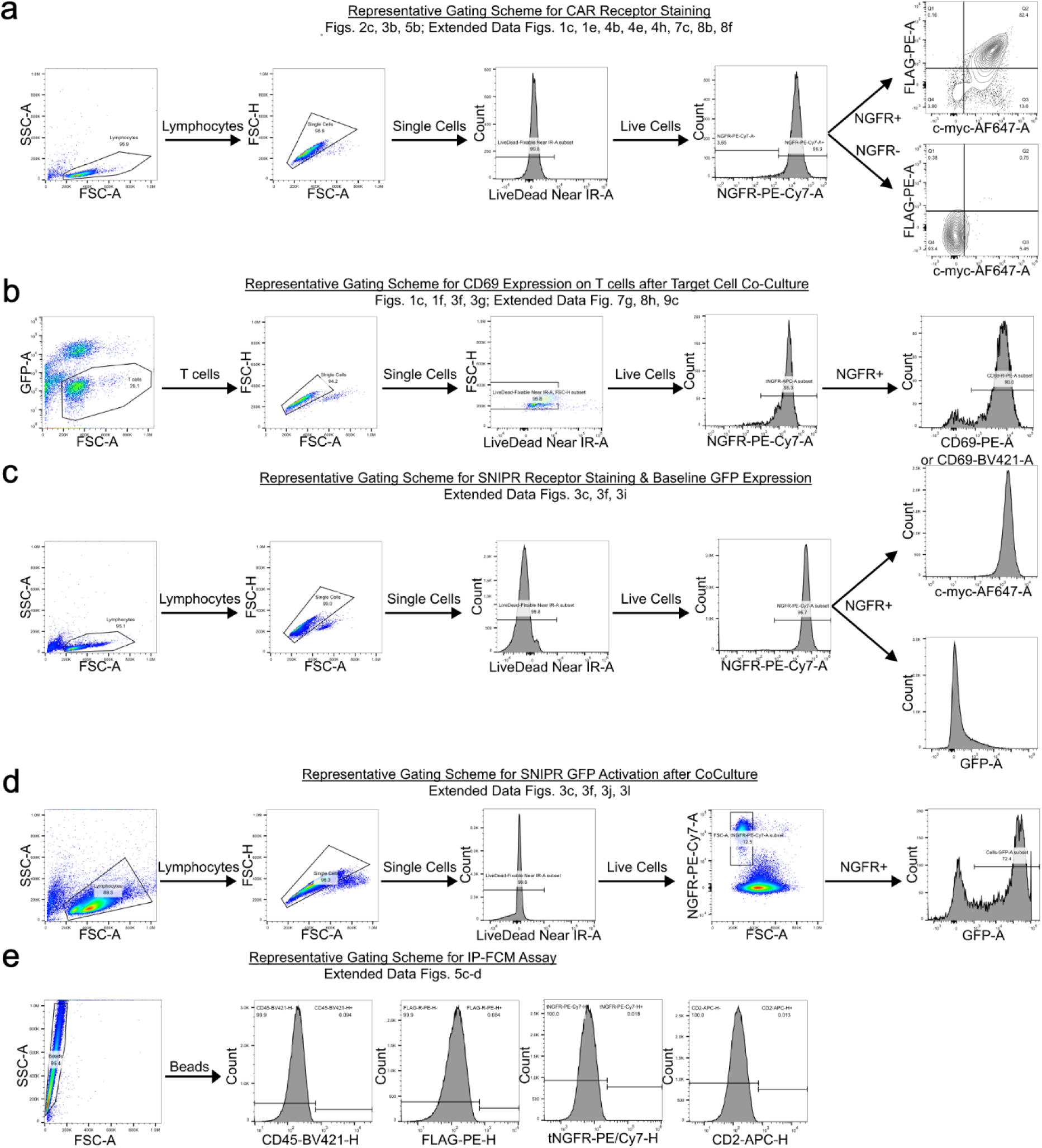
**(a-e)** Flow cytometry gating schemes for various *in vitro* experiments.

**Supplementary Figure S2.**
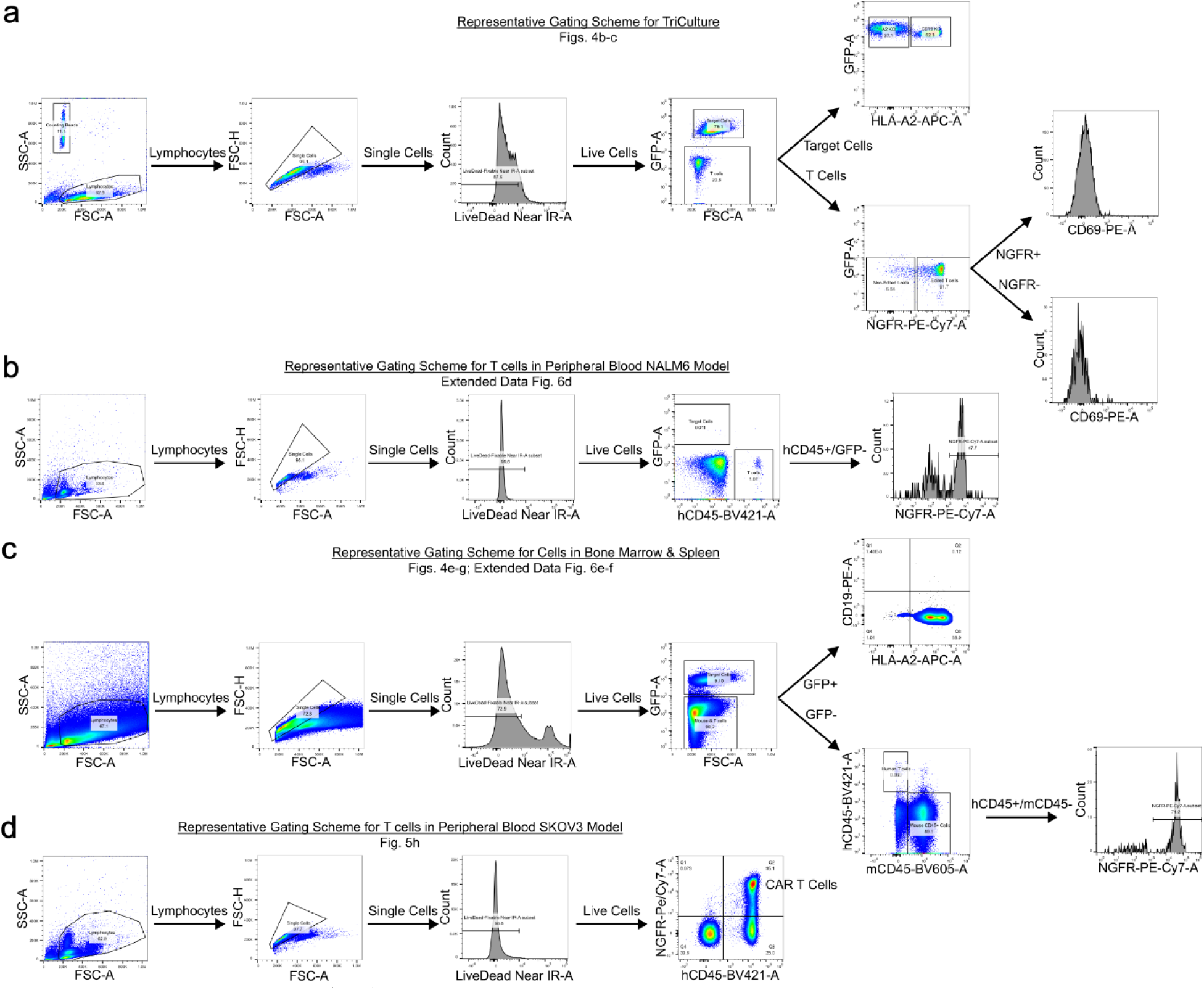
**(a-d)** Flow cytometry gating schemes for mixed co-culture and *in vivo* experiments.

